# Human Neocortical Neurosolver (HNN): A new software tool for interpreting the cellular and network origin of human MEG/EEG data

**DOI:** 10.1101/740597

**Authors:** Samuel A Neymotin, Dylan S Daniels, Blake Caldwell, Robert A McDougal, Nicholas T Carnevale, Mainak Jas, Christopher I Moore, Michael L Hines, Matti Hämäläinen, Stephanie R Jones

**Affiliations:** Dept. Neuroscience and Carney Institute for Brain Sciences, Brown University, Providence RI; Center for Biomedical Imaging and Neuromodulation, Nathan S. Kline Institute for Psychiatric Research, Orangeburg, NY; Athinoula A. Martinos Center for Biomedical Imaging, Massachusetts General Hospital, Charlestown, MA; Harvard Medical School, Boston, MA; Center for Neurorestoration and Neurotechnology, Providence VAMC; Dept. Neuroscience, Yale University, New Haven, CT; Dept. of Biostatistics, Yale University, New Haven CT

## Abstract

Magneto- and electro-encephalography (MEG/EEG) non-invasively record human brain activity with millisecond resolution providing reliable markers of healthy and disease states. Relating these macroscopic signals to underlying cellular- and circuit-level generators is a limitation that constrains using MEG/EEG to reveal novel principles of information processing or to translate findings into new therapies for neuropathology. To address this problem, we built Human Neocortical Neurosolver (HNN, https://hnn.brown.edu) software. HNN has a graphical user interface designed to help researchers and clinicians interpret the neural origins of MEG/EEG. HNN’s core is a neocortical circuit model that accounts for biophysical origins of electrical currents generating MEG/EEG. Data can be directly compared to simulated signals and parameters easily manipulated to develop/test hypotheses on a signal’s origin. Tutorials teach users to simulate commonly measured signals, including event related potentials and brain rhythms. HNN’s ability to associate signals across scales makes it a unique tool for translational neuroscience research.

## Introduction

Modern neuroscience is in the midst of a revolution in understanding the cellular and genetic substrates of healthy brain dynamics and disease due to advances in cellular- and circuit-level approaches in animal models, e.g., two-photon imaging and optogenetics. However, the translation of new discoveries to human neuroscience is significantly lacking (Badre, Frank, & Moore, 2015; Sahin et al., 2018). To understand human disease, and more generally the human condition, we must study humans. To date, EEG and MEG are the only noninvasive methods to study electrical neural activity in humans with fine temporal resolution. Despite the fact that EEG/MEG provide biomarkers of almost all healthy and abnormal brain dynamics, these so called “macro-scale” techniques suffer from difficulty in interpretability in terms of the underlying cellular- and circuit-level events. As such, there is a need for a translator that can bridge the “micro-scale” animal data with the “macro-scale” human recordings in a principled way. This is the ideal problem for computational neural modeling, where the model can have specificity at different scales.

To address this need, we developed the Human Neocortical Neurosolver (HNN), a modeling tool designed to provide researchers and clinicians an easy-to-use software platform to develop and test hypotheses regarding the neural origin of their data. The foundation of the HNN software is a neocortical model that accounts for the biophysical origin of macroscale extracranial EEG/MEG recordings with enough detail to translate to the underlying cellular- and network-level activity. HNN’s graphical user interface (GUI) provides users with an interactive tool to interpret the neural underpinnings of EEG/MEG data and changes in these signals with behavior or neuropathology.

HNN’s underlying model represents a canonical neocortical circuit based on generalizable features of cortical circuitry, with individual pyramidal neurons and interneurons arranged across the cortical layers, and layer-specific input pathways that relay spiking information from other parts of the brain, which are not explicitly modeled. Based on known electromagnetic biophysics underlying macroscale EEG/MEG signals (Jones, 2015), the elementary current generators of EEG/MEG (current dipoles) are simulated from the intracellular current flow in the long and spatially-aligned pyramidal neuron dendrites (Hämäläinen, Hari, Ilmoniemi, Knuutila, & Lounasmaa, 1993; Ikeda, Wang, & Okada, 2005; Jones, 2015; Murakami, Hirose, & Okada, 2003; Murakami & Okada, 2006; Okada, Wu, & Kyuhou, 1997). This unique construction produces equal units between the model output and source-localized data (nanoampere-meters, nAm) allowing one-to-one comparison between model and data to guide interpretation.

The extracranial macroscale nature of EEG/MEG limits the space of signals that are typically observed and studied. The majority of studies focus on quantification of event related potentials (ERPs) and low-frequency brain rhythms (<100Hz), and there are commonalities in these signals across tasks and species (Buzsáki, Logothetis, & Singer, 2013; Shin, Law, Tsutsui, Moore, & Jones, 2017). HNN’s underlying mathematical model has been successfully applied to interpret the mechanisms and meaning of these common signals, including sensory evoked responses and oscillations in the alpha (7-14 Hz), beta (15-29Hz) and gamma bands (30-80Hz) (Jones et al., 2009; Jones, Pritchett, Stufflebeam, Hämäläinen, & Moore, 2007; Lee & Jones, 2013; Sherman et al., 2016; Ziegler et al., 2010), and changes with perception (Jones et al., 2007) and aging (Ziegler et al., 2010). The model has also been used to study the impact of non-invasive brain stimulation on circuit dynamics measured with EEG (Sliva et al., 2018), and to constrain more reduced “neural mass models” of laminar activity (Pinotsis et al., 2017). In the clinical domain, HNN’s model has also been applied to study MEG-measured circuit deficits in Autism (Khan et al., 2015).

Despite these examples of use, the complexity of the original model and code hindered use by the general community. The innovation in the new HNN software is the construction of an intuitive graphical user interface to interact with the model without any coding. We offer several free and publicly-available resources to assist the broad EEG/MEG community in using the software and applying the model to their studies. These resources include an example workflow and several tutorials to study ERPs and oscillations, based on the prior studies cited above, and community sharing resources.

HNN’s GUI is designed so that researchers can simultaneously view the model’s net current dipole output and microscale features (including layer-specific responses, individual cell spiking activity, and somatic voltages) in both the time and frequency domains. HNN is constructed to be a hypothesis development and testing tool to produce circuit-level predictions that can then be directly tested and informed by invasive recordings and/or other imaging modalities. This level of scalability provides a unique tool for translational neuroscience research.

In this paper, we outline biophysiological and physiological background information that is the basis of the development of HNN, give an overview of tutorials and available data and parameter sets to simulate ERPs and low-frequency oscillations in the alpha, beta, and gamma range, and describe current distribution and online resources (https://hnn.brown.edu). We discuss the differences between HNN and other EEG/MEG modeling software packages, as well as limitations and future directions.

## Results

### Background information on the generation of EEG/MEG signals and uniqueness of HNN

#### Primary currents and the relation to forward and inverse modeling

Extracranial “macroscale” EEG/MEG are generated by large electrical currents in the brain known as primary currents **J**^p^ (Figure 1). HNN is designed to bridge the “macroscale” recordings to the underlying cellular- and circuit-level activity based on the biophysical origin of the primary electrical currents, which are assumed to be generated by the post-synaptic, intracellular longitudinal current flow in the long and spatially-aligned dendrites of a large population of synchronously-activated neocortical pyramidal neurons (Hämäläinen et al., 1993; Ikeda et al., 2005; Jones, 2015; Murakami et al., 2003; Murakami & Okada, 2006; Okada et al., 1997). Before describing how to infer the neural origin of the primary currents with HNN, we first briefly review the process of forward and inverse modeling.

**Figure 1:**
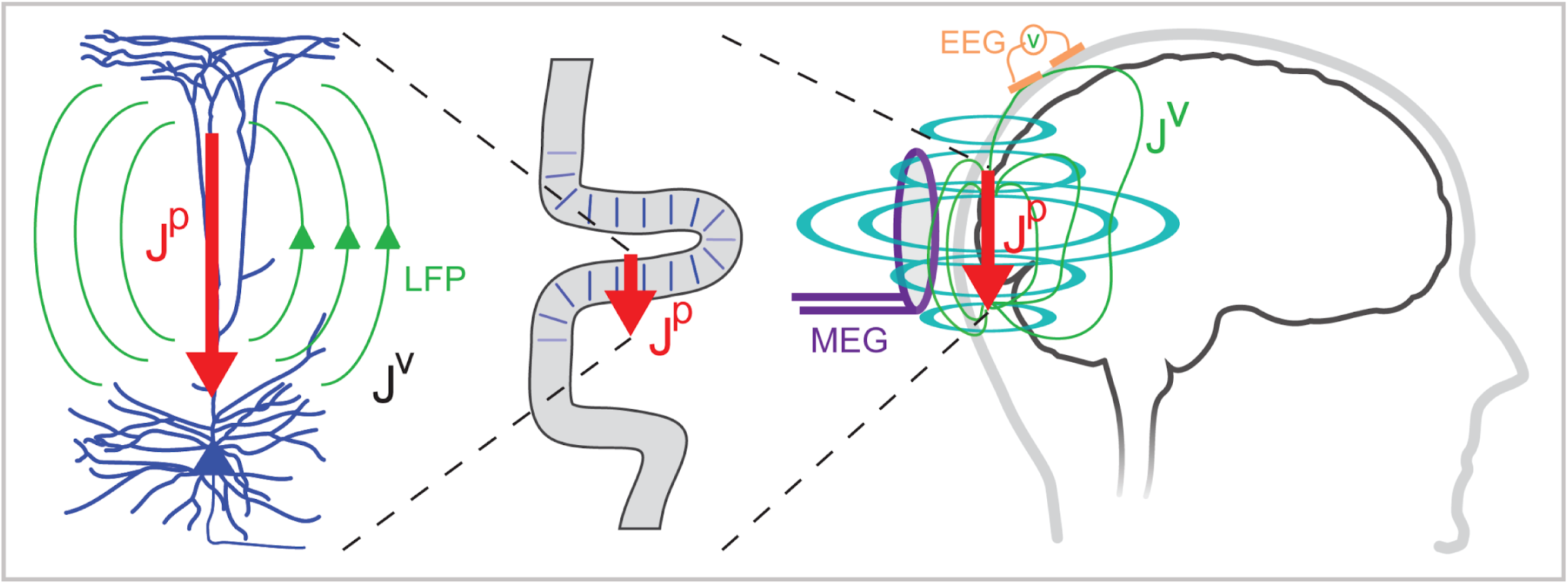
HNN bridges the “macroscale” extracranial EEG/MEG recordings to the underlying cellular- and circuit-level activity by simulating the primary electrical currents (J^P^) underlying EEG/MEG, which are generated by the postsynaptic, intracellular current flow in the long and spatially-aligned dendrites of a large population of synchronously-activated pyramidal neurons. Adapted from Jones SR, Encycl. Comput. Neurosci. 2015.

The task of computing EEG and MEG given **J**^p^ is commonly called forward modeling, governed by Maxwell’s equations. The primary currents set up a potential distribution, measurable on the scalp as EEG, and volume currents (**J**^v^) that extend through the brain tissue, the cerebrospinal fluid (CSF), the skull, and the scalp. MEG, in general, is generated by both the **J**^p^ and **J**^v^. However, in the geometry of the head, the integral effect of the volume currents to the magnetic field can be relatively easily taken into account and, therefore, modeling of MEG is in general more straightforward than the precise calculation of the electric potentials measured in EEG.

The availability of these forward models opens up the possibility to estimate the locations (**r**) and time course of the activity, **J**^p^ = **J**^p^(**r**,t), from MEG and EEG data. However, this inverse problem is fundamentally ill-posed and constraints are needed to render the problem unique. The different source localization methods, such as current dipole fitting, minimum-norm estimates, sparse source estimation methods, and beamformer approaches, differ in their capability to approximate the extent of the source activity and in their localization accuracy. However, all of these methods are capable of inferring both the location and direction of the neural currents and their time courses. Importantly, due to physiological considerations, the appropriate elementary source in all of these methods is a *current dipole*. When used in combination of geometrical models of the cortex constructed from anatomical MRI, the current direction can be related to the direction of the outer normal of the cortex: one is thus able to tell whether the estimated current is flowing outwards or inwards at a particular cortical site at a particular point in time. As such, the direction of the current flow can be related to orientation of the pyramidal neuron apical dendrites and inferred as currents flow from soma to apical tuft (up the dendrites) or apical tuft to soma (down the dendrites). There are presently several open source software packages for MEG/EEG source estimation, e.g., the MNE software, which can be employed in conjunction with HNN (Gramfort et al., 2013).

#### Inferring the neural origin of the primary currents with HNN

The focus of HNN is on the “bottom-up problem”, i.e., to study how **J**^p^ is generated by the neural circuits in the brain. Currently, the process of estimating the primary current sources (i.e., current dipoles) with inverse methods, or calculating the forward solution from **J**^p^ to the measured sensor level signal, is separate from HNN. A future direction is to integrate the top-down source estimation software with our bottom-up HNN model for all-in-one source estimation and circuit interpretation (see Discussion).

HNN’s underlying neural model contains elements that can simulate the primary current dipoles (**J**^p^) creating EEG/MEG signals in a biophysically principled manner. Specifically, HNN simulates the primary current from a canonical model of a layered neocortical column via the net intracellular electrical current flow in the pyramidal neuron dendrites in a direction parallel to the apical dendrites (see red arrow in Figures 1 and 2, and further discussion in Materials and Methods) (Hämäläinen et al., 1993; Ikeda et al., 2005; Jones, 2015; Murakami et al., 2003; Murakami & Okada, 2006; Okada et al., 1997). With this construction, the units of measure produced by the model are the same as those estimated from source localization methods, namely, nanoampere-meters (nAm), enabling one-to-one comparison of results.

**Figure 2:**
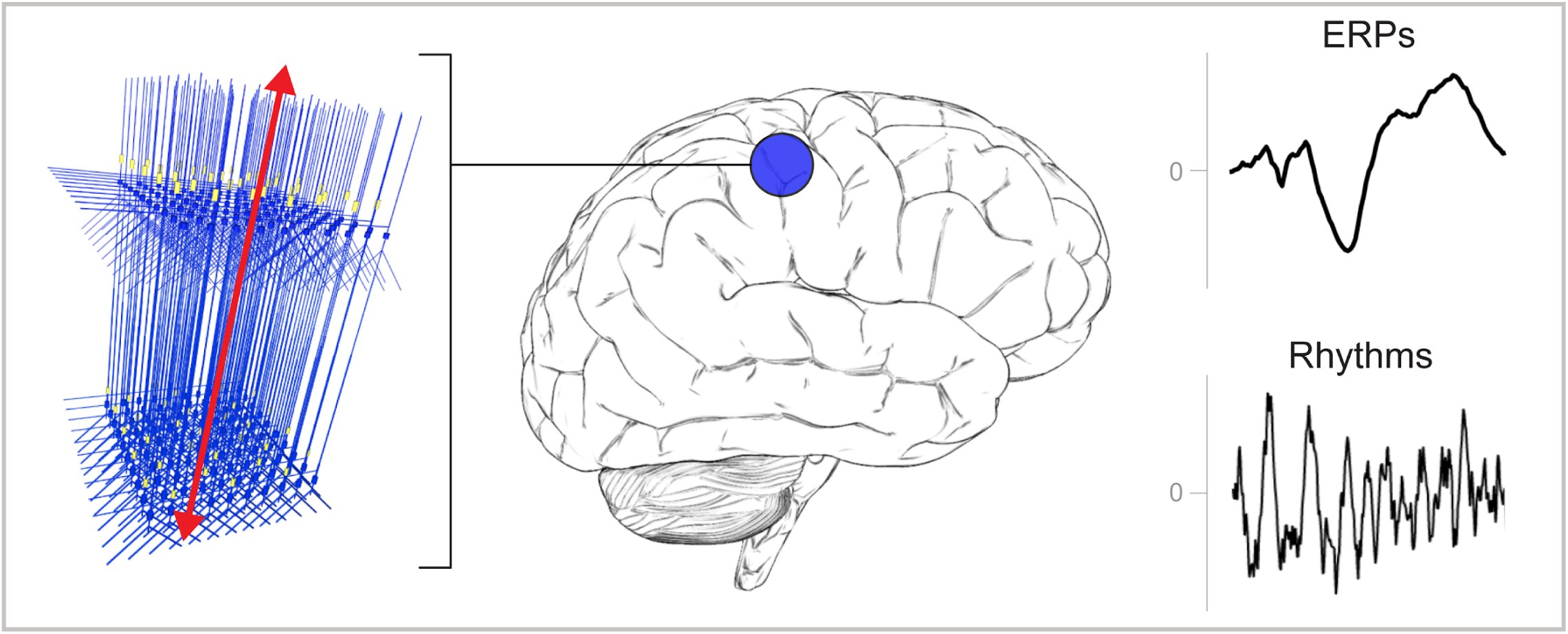
A schematic illustration of a canonical patch of neocortex that is represented by HNN’s underlying neural model. Left) 3D visualization of HNN’s model (pyramidal neurons drawn in blue, interneurons drawn in yellow). Right) Commonly measured EEG/MEG signals (ERPs and low frequency rhythms) from a single brain area that can be studied with HNN.

This construction is unique compared to other EEG/MEG modeling software (see Discussion). A necessary step in comparing model results with source localized signals is an understanding of the direction of the estimated net current in or out of the cortex, which corresponds to current flow down or up the pyramidal neuron dendrites, respectively, as discussed above. Estimation of current flow orientation at any point in time is an option in most inverse solution software that helps guide the neural interpretation, as does prior knowledge of the relay of sensory information in the cortex, see further discussion in the Tutorials part of the Results section.

By keeping model output in close agreement with the data, HNN’s underlying model has led to new and generative predictions on the origin of sensory evoked responses and low-frequency rhythms, and changes in these signals across experimental conditions (Jones et al., 2009, 2007; Khan et al., 2015; Lee & Jones, 2013; Sherman et al., 2016; Sliva et al., 2018; Ziegler et al., 2010), described further below. The macro- to micro-scale nature of the HNN software is designed to develop and test hypotheses that can be directly validated with invasive recordings or other imaging modalities (see further discussion in tutorial on Alpha and Beta rhythms).

HNN is currently constructed to dissect the cell and network contributions to signals from one source localized region of interest. Specifically, the HNN GUI is designed to simulate sensory evoked response and low-frequency brain rhythms from a single region, based on the local network dynamics and the layer-specific thalamo-cortical and cortico-cortical inputs that contribute to the local activity. As such, HNN’s underlying neocortical network represents a scalable patch of neocortex containing canonical features of neocortical circuitry (Figure 2). Ongoing expansions will include the ability to import other user-defined cell types and circuit models into HNN, as well as the ability to simulate the interactions among multiple neocortical areas (see Discussion). Of note, users can still benefit from our software if they are working with data directly from EEG/MEG sensor rather than source localized signals. The primary currents are the foundation of the sensor signal and, as such, can have similar activity profiles (e.g., compare source localized tactile evoked response in Figure 4 and sensor level response in Figure 5).

### Overview of HNN’s default canonical neocortical column template network

#### Neocortical column structure

Here, we give an overview of the main features that are important to understand in order to begin exploring the origin of macroscale evoked responses and brain rhythms, and we provide details on how these features are implemented in HNN’s template model. Further details can be found in the Materials and Methods section, in our prior publications (e.g., Jones et al., 2009), and on our website https://hnn.brown.edu.

Given that the primary electrical current that generates EEG/MEG signals comes from synchronous activity in pyramidal neuron (PN) dendrites across a large population, there are several key features of neocortical circuitry that are essential to consider when simulating these currents. While there are known differences in microscale circuitry across cortical areas and species, many features of neocortical circuits are remarkably similar. We assume these conserved features are minimally sufficient to account for the generation of evoked responses and brain rhythms measured with EEG/MEG, and we have harnessed this generalization into HNN’s foundational model, with success in simulating many of these signals using the same template model (see Introduction). These canonical features include:

**(I)** A 3-layered structure with pyramidal neurons in the supragranular and infragranular layers whose dendrites span across the layers and are synaptically coupled to inhibitory interneurons in a 3-to-1 ratio, of pyramidal to inhibitory cells (Figure 3A). Of note, cells in the granular layer are not explicitly included in the template circuit. This initial design choice was based on the fact that macroscale current dipoles are dominated by PN activity in supragranular, and infragranular layers. Thalamic input to granular layers is presumed to propagate directly to basal and oblique dendrites of PN in the supragranular and infragranular layers. In the model, the thalamic input synapses directly onto these dendrites.

**Figure 3:**
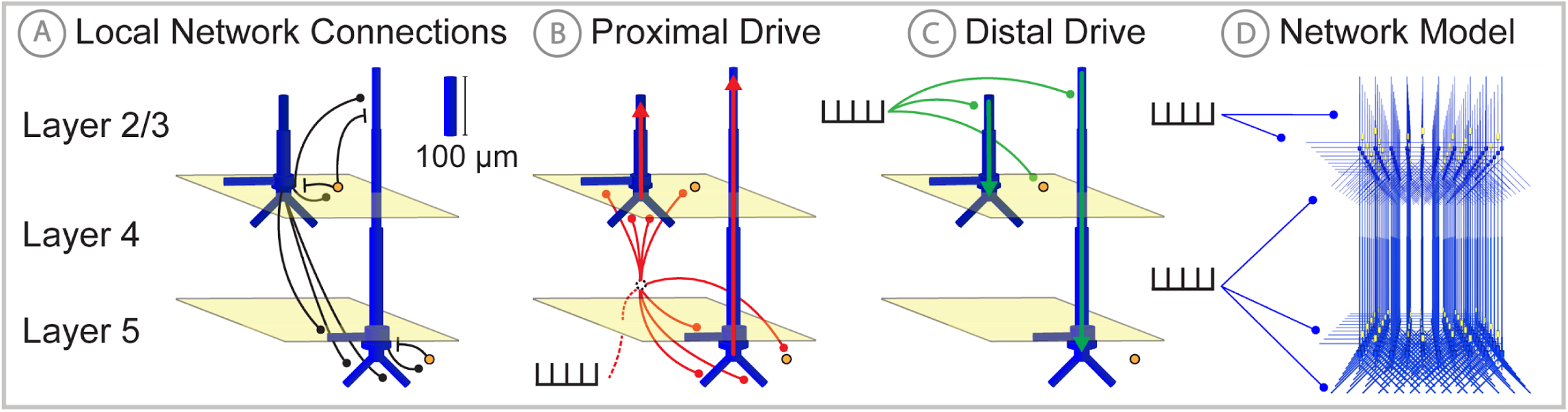
Schematic illustrations of HNN’s underlying neocortical network model. **(A)** Local Network Connectivity: GABAergic (GABAA/GABAB; lines) and glutamatergic (AMPA/NMDA; circles) synaptic connectivity between single-compartment inhibitory neurons (orange circles) and multi-compartment layer 2/3 and layer 5 pyramidal neurons (blue neurons). Excitatory to excitatory connections not shown, see Materials and Methods. **(B)** Exogenous proximal drive representing lemniscal thalamic drive to cortex. User defined trains or bursts of action potentials (see tutorials described in Results) are simulated and activate post-synaptic excitatory synapses on the basal and oblique dendrites of layer 2/3 and layer 5 pyramidal neurons as well as the somata of layer 2/3 and layer 5 interneurons. These excitatory synaptic inputs drive current flow up the dendrites towards supragranular layers (red arrows). **(D)** Exogenous distal drive representing cortical-cortical inputs or non-lemniscal thalamic drive that synapses directly into the supragranular layer. User defined trains of action potentials are simulated and activate post-synaptic excitatory synapses on the distal apical dendrites of layer 5 and layer 2/3 pyramidal neurons as well as the somata of layer 2/3 interneurons. These excitatory synaptic inputs push the current flow down towards the infragranular layers (green arrows). **(D)** The full network contains a scalable number of pyramidal neurons in layer 2/3 and layer 5 in a 3-to-1 ratio with inhibitory interneurons, activated by user defined layer specific proximal and distal drive (see Materials and Methods for full details).

The number of cells in the network is adjustable in the Local Network Parameters window via the Cells tab, while maintaining at 3-to-1 pyramidal to inhibitory interneuron ratio in each layer. The connectivity pattern is fixed, but the synaptic weights between cell types can be adjusted in the Local Network menu and the Synaptic Gains menu. Macroscale EEG/MEG signals are generated by the synchronous activity in large populations of PN neurons. Evoked responses are typically on the order of 10-100nAm, and are estimated to be generated by the synchronous spiking activity of the order of tens of thousands of pyramidal neurons. Low-frequency oscillations are larger in magnitude and are on the order of 100-1000nAm, and are estimated to be generated by the subthreshold activity of on the order of a million pyramidal neurons (Jones et al., 2009, 2007; Murakami & Okada, 2006) While HNN is constructed with the ability to adjust local network size, the magnitude of these signals can also be conveniently matched by applying a scaling factor to the model output, providing an estimate on the number of neurons that contributed to the signal.

**(II)** Exogenous driving input through two known layer-specific pathways. One type of input represents excitatory synaptic drive that comes from the lemniscal thalamus and contacts the cortex in the granular layers, which then propagates to the proximal PN dendrites in the supragranular and infragranular layers and somata of the inhibitory neurons; this input is referred to as proximal drive (Figure 3B). The other input represents excitatory synaptic drive from higher-order cortex or non-specific thalamic nuclei that synapses directly into the supragranular layers and contacts the distal PN dendrites and somata of the inhibitory neuron; this input is referred to as distal drive (Figure 3C). The networks that provide proximal and distal input to the local circuit (e.g., thalamus and higher order cortex) are not explicitly modelled, but rather these inputs are represented by simulated trains of action potentials that activate excitatory post-synaptic receptors in the local network. The temporal profile of these action potentials is adjustable depending on the simulation experiment and can be represented as single spikes, bursts of input, or rhythmic bursts of input. There are several ways to change the pattern of action potential drive through different buttons built into the HNN GUI: Evoked Inputs, Rhythmic Proximal Inputs and Rhythmic Distal Inputs. The dialog boxes that open with these buttons allow creation and adjustment of patterns of evoked response drive or rhythmic drive to the network (see tutorials described in Results section for further details).

**(III)** Exogenous drive to the network can also be generated as excitatory synaptic drives following a Poisson process to the somata of chosen cell classes or as tonic input simulated as a somatic current clamp with a fixed current injection. The timing and duration of these drives is adjustable.

Further details of the biophysics and morphology of the cells and architecture of the local synaptic connectivity profiles in the template network can be found in the Materials and Methods section. As the use of our software grows, we anticipate other cells and network configurations will be made available as template models to work with via open source sharing (see Discussion).

#### Parameter tuning in HNN’s template network model

HNN’s template model is a large-scale model simulated with thousands of differential equations and parameters, making the parameter optimization process challenging. The process for tuning this canonical model and constraining the space of parameters to investigate the origin of ERPs and low-frequency oscillations was as follows. First, the individual cell morphologies and physiologies were constrained so individual cells produced realistic spiking patterns to somatic injected current (detailed in Methods). Second, the local connectivity within and among cortical layers was constructed based on a large body of literature from animal studies (detailed in Methods). All of these equations and parameters were then fixed, and the only parameters that were originally tuned to simulate ERPs and oscillations were the timing and the strength of the exogenous drive to the local network. This drive represented our “simulation experiment” and was based on our hypotheses on the origin of these signals motivated by literature and on matching model output to features of the data (see tutorials described in Results). The HNN GUI was constructed assuming ERPs and low-frequency oscillations depend on layer-specific exogenous drives to the network. The simulation experiment workflow and tutorials described below are in large part based on “activating the network” by defining the characteristics of this layer-specific drive. Default parameter sets are provided as a starting point from which the underlying parameters can be interactively manipulated using the GUI, and additional exogenous driving inputs can be created or removed.

Automated parameter optimization is also available in HNN and is specifically designed to accurately reproduce features of an ERP waveform based on the temporal spacing and strength of the exogenous driving inputs assumed to generate the ERP. Before taking advantage of HNN’s automated parameter optimization, we strongly encourage users to begin by understanding our ERP tutorial, and hand-tuning parameters using one of our default parameter sets to get an initial representation of the recorded data. The identification of an appropriate number of driving inputs and their approximate timings and strengths serves as a starting point for the optimization procedure (described in the ERP Model Optimization section below). Hand tuning of parameters and visualizing the resultant changes in the GUI will enable users to understand how specific parameter changes impact features of the current dipole waveform.

Importantly, the biophysical constraints on the origin of the current dipoles signal (discussed above) will dictate the output of the model and necessarily limits the space of parameter adjustments that can accurately account for the recorded data. The same principle underlies the fact that a limited space of signals are typically studied at the macroscale (ERPs and low frequency oscillations). A parameter sensitivity analysis on perturbations around the default ERP parameter sets confirmed that a subset of the parameters have the strongest influence on features of the ERP waveform (see Supplementary Materials). Insights from GUI interactive hand tuning and sensitivity analyses can help narrow the number of parameters to include in the subsequent optimization procedure and greatly decrease the number of simulations required for optimization.

### HNN GUI overview and interactive simulation experiment workflow

The HNN GUI is designed to allow researchers to link macro-scale EEG/MEG recordings to the underlying cellular- and network-level generators. Currently available visualizations include, a direct comparison of simulated electrical sources to recorded data with calculated goodness of fit estimates, layer-specific current dipole activity, individual cell spiking activity, and individual cell somatic voltages (Figure 4B-D). Results can be visualized in both the time and frequency domain. Based on its biophysically detailed design, the output of HNN’s model and recorded source-localized data have the same units of measure, nAm. By closely matching the output of the model to recorded data in an interactive manner, users can test and develop hypotheses on the cell and network origin of their signals.

**Figure 4:**
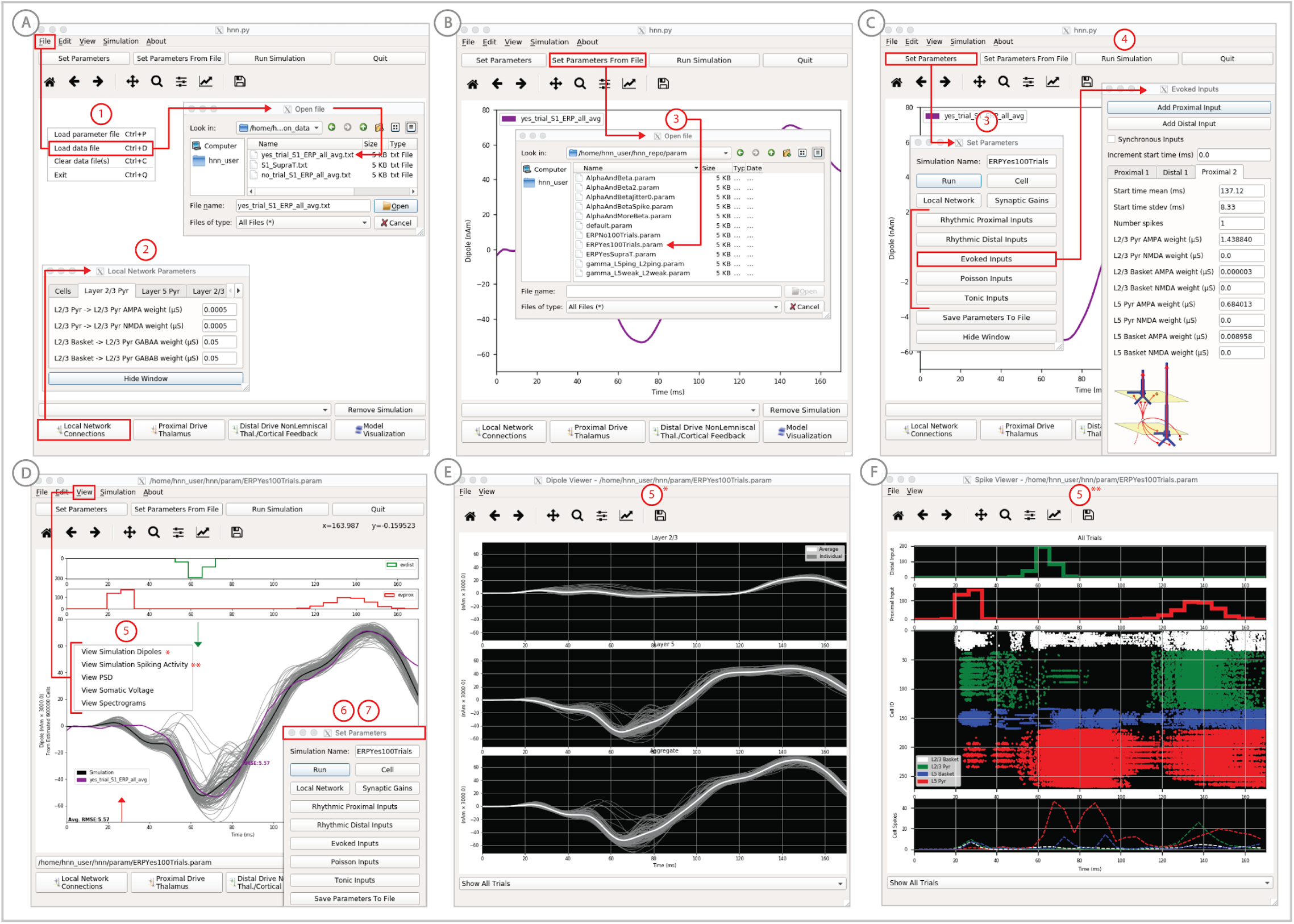
An example workflow showing how HNN can be used to link the macroscale current dipole signal to the underlying cell and circuit activity. The example shown is for a perceptual threshold level tactile evoked response (50% detected) from SI (Jones et al 2007; see ERP Tutorial text for details). **(A)** Steps 1 and 2: load data and define the local network structure. **(B)** Step 3: activate the local network, starting with a predefined parameter set; shown here for the parameter set for perceptual threshold-level evoked response (ERPYes100Trials.param) **(C)** Step 3 and 4: adjust the evoked input parameters according to user defined hypotheses and simulation experiment, and run the simulation. **(D)** Step 5: visualize model output; the net current dipole will be displayed in the main GUI window and microcircuit details, including layer-specific responses, cell membrane voltages, and spiking profiles (**E and F**) are shown by choosing them from the View pull down menu. Parameters can be adjusted to hypothesized circuit changes under different experimental conditions (e.g. see Figure 5).

The process for simulating evoked responses or brain rhythms from a single region of interest is to first define the network structure, and then to “activate” the network with exogenous driving input based on your hypotheses and simulation experiment. HNN’s template model provides the initial network structure. The choice of “activation” to the network depends on the simulation experiment. The GUI design is motivated by our prior published studies and was built specifically to simulate sensory evoked responses or spontaneous rhythms, or a combination of the two (Jones et al., 2009, 2007; Khan et al., 2015; Lee & Jones, 2013; Sherman et al., 2016; Sliva et al., 2018; Ziegler et al., 2010). The tutorials described in the Results section below details examples of how to “activate” the network to simulate sensory evoked responses and spontaneous rhythms. Here, we outline a typical simulation experiment workflow.

In practice, users apply the following interactive workflow, as in Figure 4 and detailed further in the tutorials with an example tactile evoked response from somatosensory cortex (data from Jones et al., 2007).

**(Step 1)** Load EEG/MEG data (*blue*). (Step 1 is optional.)

**(Step 2)** Define the cortical column network structure. The default template network is automatically loaded when HNN starts. Default parameters describing the local network can be adjusted by clicking the Set Parameters button on the GUI and then Local Network Parameters, or directly from the Local Network Parameters button on the GUI (Figure 4A).

**(Step 3)** “Activate” the local network by defining layer-specific, exogenous driving inputs (Figure 3 B,C). The drive represents input to the local circuit from thalamus and/or other cortical areas and can be in the form of (i) spike trains (single spikes or bursts of rhythmic input) that activate post-synaptic targets in the local network, (ii) current clamps (tonic drive), or (iii) noisy (Poisson) synaptic drive. The choice of input parameters depends on your hypotheses and “simulation experiment”. In the example simulation, predefined evoked response parameters where loaded in via the Set Parameters From File button and choosing the file “ERPYes100Trials.param”; this is also the default evoked response parameter set loaded when starting HNN (Figure 4B). The Evoked Input parameters are then viewed in the Set Parameters dialog box under Evoked Inputs (Figure 4C). The Evoked Inputs parameters are described further in the tutorials below.

**(Step 4)** Run simulation and directly compare model output (*black*) and data (*purple*) with goodness of fit calculations (root mean squared error, RMSE, between data and averaged simulation) (Figure 4D).

**(Step 5)** Visualize microcircuit details, including layer-specific responses, cell membrane voltages, and spiking profiles by choosing from the View pull down menu (Figure 4D, E, F).

**(Step 6)** Adjust parameters through the Set Parameters dialog box to develop and test predictions on the circuit mechanisms that provide the best fit to the data. With any parameter adjustment, the change in the dipole signal can be viewed and compared with the prior simulation to infer how specific parameters impact the current dipole waveform. Prior simulations can be maintained in the GUI or removed. For ERPs, automatic parameter optimization can be iteratively applied to tune the parameters of the exogenous driving inputs to find those that provide the best initial fit between the simulated dipole waveform and the EEG/MEG data (see further details below).

**(Step 7)** To infer circuit differences across experimental conditions, once a fit to one condition is found, adjustments to relevant cell and network parameters can be made (guided by user-defined hypotheses), and the simulation can be re-run to see if predicted changes account for the observed differences in the data. A list of the GUI-adjustable parameters in the model can be found in the “Tour of the GUI” section of the tutorials on our website. HNN’s GUI was designed so that users could easily find the adjustable parameters from buttons and pull down menus on the main GUI leading to dialogue boxes with explanatory labels.

As a specific example on how to use HNN as a hypothesis testing tool, we have used HNN to evaluate hypothesized changes in EEG measured neural circuit dynamics with non-invasive brain stimulation (Figure 5). We measured somatosensory evoked responses from brief threshold-level taps to the middle finger tip before and after 10 minutes of ∼10Hz transcranial alternating current stimulation (tACS) over contralateral somatosensory cortex (see Sliva et al., 2018 for details). The magnitude of an early peak near ∼70ms in the tactile evoked response increased after the tACS session (Figure 5, top left). Based on prior literature, we hypothesized that the observed difference was due to changes in synaptic efficacy in the local network induced by the tACS (Kronberg, Bridi, Abel, Bikson, & Parra, 2017; Rahman, Lafon, Parra, & Bikson, 2017). To test this hypothesis, we first used HNN to simulate the pre-tACS evoked responses, following the evoked response tutorial in our software (see Tutorial below). Once the pre-tACS condition was accounted for, we then adjusted the synaptic gain between the excitatory and inhibitory cells in the network using the HNN GUI and re-simulated the tactile evoked responses. We tested several possible gain changes between the populations. HNN showed that a two-fold increase in synaptic strength of the inhibitory connections, as opposed to an increase in the excitatory connections or in total synaptic efficacy, could best account for the observed differences in the data (compare blue in red curves in Figure 5). By viewing the cell spiking profiles in each condition (Figure 5, bottom right), HNN further predicted that the increase in the magnitude of the ∼70ms peak coincided with increased firing in the inhibitory neuron population and decreased firing in the excitatory pyramidal neurons in the post-tACS compared to the pre-tACS window. These detailed predictions can guide further experiments and follow-up testing in animal models or with other human imaging experiments. Follow up testing of model derived predictions is described further in the alpha/beta tutorial below.

**Figure 5:**
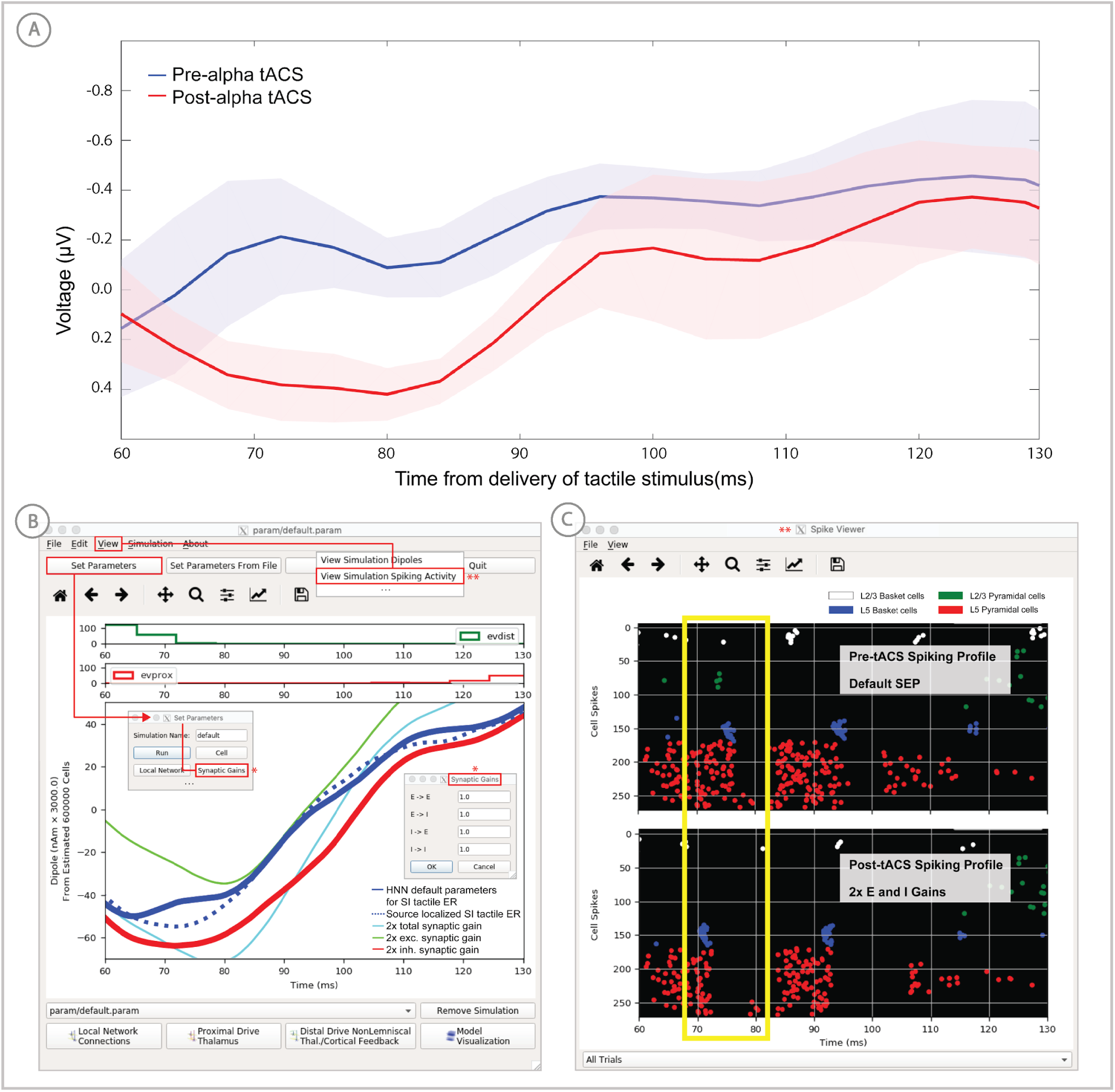
Application of HNN to test alternative hypotheses on the circuit level impact of tACS on the somatosensory tactile evoked response (adapted from Sliva et al 2018). **(A)** The early tactile evoked response from above somatosensory cortex before and after 10 minutes of 10Hz alternating current stimulation over SI shows that the ∼70ms peak is more prominent in the post-tACS condition. Note that the timing of this peak in the sensor level signal is analogous to the 70ms peak in the source localized signal in Figure 4B, since the tactile stimulation was the same in both studies and the early signal from SI is similar both at the source and sensor level. **(B)** HNN was applied to investigate the impact of several possible tACS induced changes in local synaptic efficacy and identify which could account for the observed evoked response data. The parameters in HNN were first adjusted to account for the pre-tACS response using the default HNN parameter set (solid blue line). The synaptic gains between the different cell types was then adjusted through the Set Parameters dialog box to predict that 2x gain in the local inhibitory synaptic weights best accounted for the post-tACS evoked response. (**C)** Simultaneous viewing of the cell spiking activity further predicted that there is less pyramidal neuron spiking at 70ms post-tACS, despite the more prominent 70ms current dipole peak.

### Tutorials on ERPs and low-frequency oscillations

HNN’s tutorials are designed to teach users how to simulate the most commonly studied EEG/MEG signals, including sensory evoked responses and low-frequency oscillations (alpha, beta, and gamma rhythms) by walking users through the workflow we applied in our prior studies of these signals. The data and parameter sets used in these studies are distributed with the software, and the interactive GUI design was motivated by this workflow. In completing each tutorial, users will have a sense of the basic structure of the GUI and the process for manipulating relevant parameters and viewing results. From there, users can begin to develop and test hypotheses on the origin of their own data. Below we give a basic overview of each tutorial. The HNN website (https://hnn.brown.edu) provides additional information and example exercises for further exploration.

#### Sensory evoked responses

We have applied HNN to study the neural origin of tactile evoked responses localized with inverse methods to primary somatosensory cortex from MEG data (Jones et al., 2007). In this study, the tactile evoked response was elicited from a brief perceptual threshold level tap - stimulus strength maintained at 50% detection -- to the contralateral middle finger tip during a tactile detection experiment (experimental details in Jones et al., 2007, 2009). The average tactile evoked response during detected trials is shown in Figure 4. The data from this study is distributed with HNN installation.

Following the workflow described above, the process for reproducing these results in HNN is as follows.

**Steps 1 & 2**: Load the evoked response data distributed with HNN, “yes_trial_SI_ERP_all_avg.txt”. The data shown in Figure 4B will be displayed. Adjust parameters defining the automatically loaded default local network, if desired.

**Step 3:** “Activate” the local network. In prior publications, we showed that this tactile evoked response could be reproduced in HNN by “activating” the network with a sequence of layer-specific proximal and distal spike train drive to the local network, which is distributed with HNN in the file “ERPYes100Trials.param”.

The sequence described below was motivated by intracranial recordings in non-human primates, which guided the initial hypothesis testing in the model. Additionally, we established with inverse methods that at the prominent ∼70ms negative peak (Figure 4D), the orientation of the current was into the cortex (e.g. down the pyramidal neuron dendrites), consistent with prior intracranial recordings (see Jones et al., 2007). As such, in this example, negative current dipole values correspond to current flow down the dendrites, and positive values up the dendrites. In sensory cortex, the earliest evoked response peak corresponds to excitatory synaptic input from the lemniscal thalamus that leads to current flow out of the cortex (e.g. up the dendrites). This earliest evoked response in somatosensory cortex occurs at ∼25ms. The corresponding current dipole positive peak is small for the threshold tactile response in Figure 4D, but clearly visible in Figure 10 for a suprathreshold (100% detection) level tactile response.

The drive sequence that accurately reproduced the tactile evoked response consisted of “feedforward” / proximal input at ∼25 ms post stimulus, followed by “feedback” / distal input at ∼60 ms, followed by a subsequent “feedforward” / proximal input at ∼125 ms (Gaussian distribution of input times on each simulated trial, Figure 4C). This “activation” of the network generated spiking activity and a pattern of intracellular dendritic current flow in the pyramidal neuron dendrites in the local network to reproduce the current dipole waveform, many features of which fell naturally out of the local network dynamics (details in Jones et al., 2007). This sequence can be interpreted as initial “feedforward” input from the lemniscal thalamus followed by “feedback” input from higher-order cortex or non-lemniscal thalamus, followed by a re-emergent leminsical thalamic drive. A similar sequence of information flow likely applies to most sensory evoked signals. The inputs are distinguished with red and green arrows (corresponding to proximal and distal input, respectively) in the main GUI window. The number, timing, and strength (post-synaptic conductance) of the driving spikes were manually adjusted in the model until a close representation of the data was found (see section on parameter tuning above). To account for some variability across trials, the exact time of the driving spikes for each input was chosen from a Gaussian distribution with a mean and standard deviation (see Evoked Inputs dialog box, Figure 4C, and green and red histograms on the top of the GUI in Figure 4D). The gray curves in Figure 4D show 25 trials of the simulation (decreased from 100 trials in the Set Parameters, Run dialog box) and the black curve is the average across simulations. The top of the GUI windows displays histograms of the temporal profile of the spiking activity providing the sequence of proximal (red) and distal (green) synaptic input to the local network across the 25 trials. Note, a scaling factor was applied to net dipole output to match to the magnitude of the recorded ERP data and used to predict the number of neurons contributing to the recorded ERP. This scaling factor is chosen from Set Parameters, Run dialog box, and is shown as 3000 on the y-axis of the main GUI window in Figure 4D. Note that the scaling factor is used to predict the number of pyramidal neurons contributing to the observed signal. In this case, since there are 100 pyramidal neurons in each of layers 2/3 and 5, that amounts to 600,000 neurons (200 neurons x 3000 scaling factor) contributing to the evoked response, consistent with the experimental literature (described in Jones et al., 2007, 2009).

Based on the assumption that sensory evoked responses will be generated by a layer-specific sequence of drive to the local network similar to that described above, HNN’s GUI was designed for users to begin simulating evoked responses by starting with the aforementioned default sequence of drive that is defined when starting HNN and by loading in the parameter set from the “ERPYes100Trials.param” file, as described above. The Evoked Inputs dialog box (Figure 4C) shows the parameters of the proximal and distal drive (number, timing, and strength) used to produce the evoked response in Figure 4D. Here, there were two proximal drives and one distal drive to the network. These parameters were found by first hand tuning the inputs to get a close representation of the data and then running the parameter optimization procedure described below.

**Step 4** The evoked response shown in Figure 4 is reproduced by clicking the “Run Simulation” button at the top of the GUI, and the RMSE of the goodness of fit to the data is automatically calculated and displayed. Additional network features can also be visualized through pull down menus (**Step 5**).

Evoked response parameters can now be adjusted, and additional inputs can be created or removed to account for the user-defined “simulation experiment” and hypothesis testing goals (**Step 6**). With each parameter change, a new parameter file will be saved by renaming the simulation under “Simulation Name” in the “Set Parameters” dialog box (see Figure 4C). From here, other cell or network parameters can be adjusted to compare across conditions (**Step 7).**

#### Alpha and beta rhythms

We have applied HNN to study the neural origin of spontaneous rhythms localized to the primary somatosensory cortex from MEG data; it is often referred to as the mu-rhythm, and it contains a complex of (7-14Hz) alpha and (15-29Hz) beta frequency components (Jones et al., 2009). A 1-second time frequency spectrogram of the spontaneous unaveraged SI rhythm from this study is shown in Figure 6. This data is distributed on the HNN website (“SI_ongoing.txt”), and contains 1000 1-second epochs of spontaneous data (100 trials each from 10 subjects). The data is plotted in HNN through the “View → View Spectrograms” menu item, followed by “Load Data” and then selecting the “SI_ongoing.txt” file. Note that it may take a few minutes to calculate the wavelet transforms for all 1000 1-second trials included. Next, select an individual trial (e.g. trial 32) from the drop-down menu. The dipole waveform from a single 1-second epoch will then be shown in the top and the corresponding time-frequency spectrogram. The spectrogram is automatically calculated and displayed at the bottom, as seen in Figure 6. Notice that this rhythm contains brief bouts of alpha or beta activity that will occur at different times in different trials due to the spontaneous, non-stationary nature of the signals. As such, when averaged across trials, bands of alpha and beta activity appear continuous in the spectrogram (data not shown, see Jones et al., 2007; Jones, 2016) and will create peaks at alpha and beta in a power spectral density (Figure 8C). Since the alpha and beta components of this rhythm are not time locked across trials, it is difficult to directly compare the waveform of the recorded data with model output. Rather, to assess the goodness of fit of the model, we compared features of the simulated rhythm to the data (see Jones et al., 2009), including peaks in the power spectral density, as described below. Since we can not directly compare the waveform of this rhythm with the model output, rather than first loading the data, we begin this tutorial with Step 3, “activating” the network, using the default local network defined when starting HNN.

**Figure 6:**
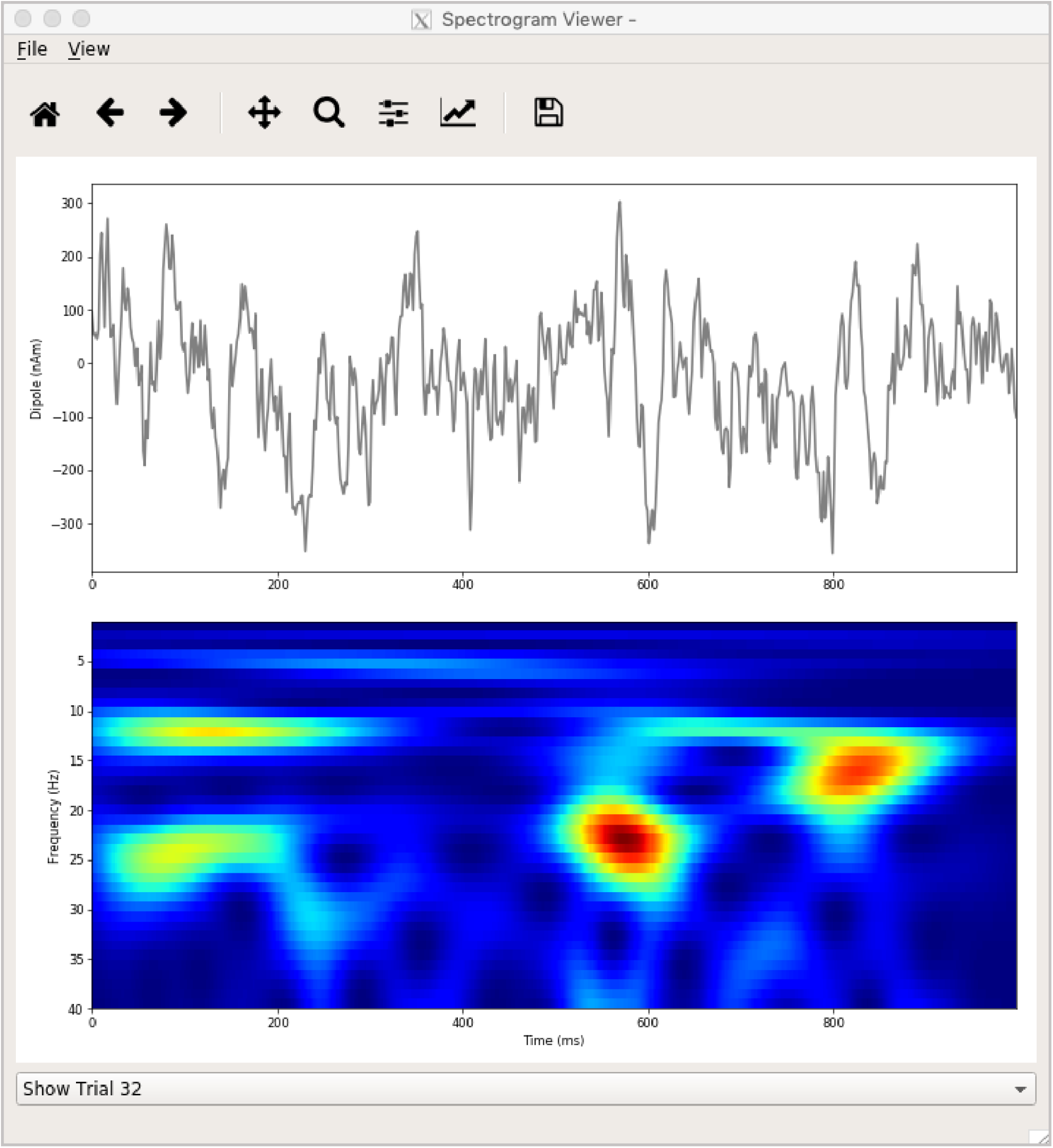
Example spontaneous data from a current dipole source in SI showing transient alpha (∼12 Hz) and beta (∼15-30 Hz) components (data as in Jones et al 2009). The data file (“SI_ongoing.txt”) used to generate these outputs is provided with HNN and plotted through the “View → View Spectrograms” menu item, followed by “Load Data”, and then selecting the file.

**Step 3:** “Activate” the local network. In prior publications, we have simulated low-frequency alpha and beta rhythms through patterns of rhythmic drive (repeated bursts of spikes) through proximal and distal projection pathways. These patterns of drive were again motivated by literature and by tuning the parameters to match features of the model output to the recorded data (see Jones et al., 2009; Sherman et al., 2016).

We begin by describing the process for simulating a pure alpha frequency rhythm only, and we then describe how a novel prediction for the origin of beta events emerged (Sherman et al. PNAS 2016). Motivated by a long history of research showing alpha rhythms in neocortex rely on ∼10Hz bursting in the thalamus, we tested the hypothesis that ∼10Hz bursts of drive through proximal and distal projection pathways (representing lemniscal and non-lemniscal thalamic drive) could reproduce an alpha rhythm in the local circuit. The burst statistics (number of spikes and inter-burst interval chosen from a Gaussian distribution), strength of the input (post-synaptic conductance), and delay between the proximal and distal input, were manually adjusted until a pure alpha rhythm sharing features of the data was found. We showed that when ∼10Hz bursts of proximal and distal drives are subthreshold and arrive to the local network in anti-phase (∼50ms delay) a pure alpha rhythm emerges (Jones et al., 2009; Ziegler et al., 2010).

The parameters of this drive are distributed with HNN in the file “Alpha.param”, loaded through the Set Parameters From File button and viewed in the Set Parameters dialog box under Rhythmic Proximal and Rhythmic Distal inputs (Figure 7A). Note that the start time mean of the ∼10Hz Rhythmic Proximal and Rhythmic Distal Inputs are delayed by 50ms. The HNN GUI in Figure 7B displays the simulated current dipole output from this drive (middle), the histogram of the proximal and distal driving spike trains (top), and the corresponding time-frequency domain response (bottom). This GUI window is automatically constructed when rhythmic inputs are given to the network, and HNN is designed to easily define rhythmic input to the network via the Set Parameters dialog box. A scaling factor was also applied to this signal (via Set Parameters, Run dialog box) and is shown as 300,000 on the y-axis of the main GUI window example in Figure 7B. The 300,000 scaling factor predicts that 60,000,000 PNs (300,000 x 200 PNs) contribute to the measured signal.

**Figure 7:**
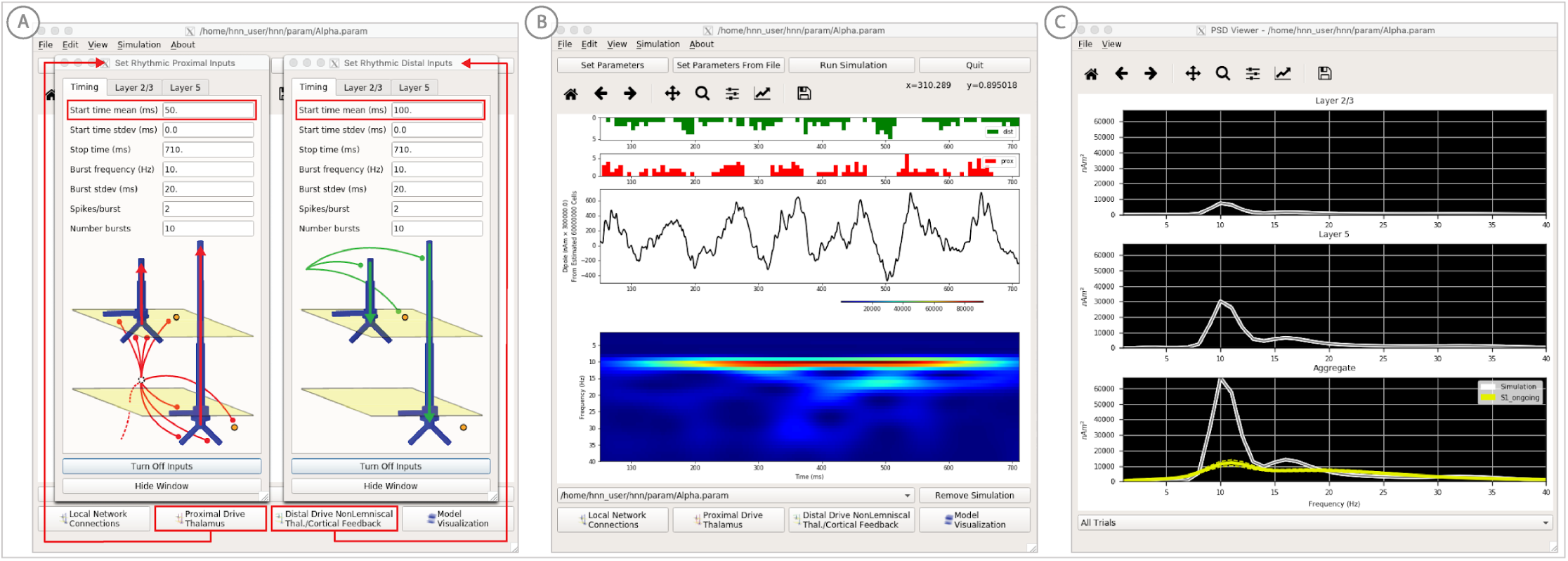
An example workflow for simulating alpha frequency rhythm (Jones et al 2009; Ziegler et al 2010; see Alpha and Beta Rhythms Tutorial text for details). **(A)** Here we are using the default HNN network configuration and not directly comparing the waveform to data, so begin with Step 3: activate the local network. Motivated by prior studies (see text), in this example alpha rhythms were simulated by driving the network with ∼10 Hz bursts (presumed to be generated by thalamus) to the local network through proximal and distal projection pathways. The parameter set describing these burst is provided in the Alpha.param file and loaded through the Set Parameters From File button. Adjustable burst drive parameter are shown and here were set with a 50ms delay between the ∼10 proximal and distal drive (red boxes). **(B)** Step 4: running the simulation with the “Run Simulation” button, shows that a continuous alpha rhythm emerged in the current dipole signal (middle dipole time trace; bottom time-frequency representation). Green and red histograms at the top display the defined distal and proximal burst drive patterns, respectively. **(C)** Step 5: additional network features, including layer specific power spectral density plots as shown can be visualized through the “View” pull down menu, and compared to data (here compared to the spontaneous SI data shown in Figure 6). Features of the burst drive can be adjusted (panel **A**) and corresponding changes in the current dipole signals studied (Steps 6 and 7, see Figure 8).

**Step 4** The alpha rhythm shown in Figure 7B is reproduced by clicking the “Run Simulation” button at the top of the GUI, Additional network features, including power-spectral density plots, can also be visualized through the pull down menus (**Step 5**).

**Steps 6 and 7:** Rhythmic input parameters can be adjusted to account for the user defined “simulation experiment” and hypothesis testing goals.

The goal in our prior study was to reproduce the alpha / beta complex of the SI mu-rhythm. By hand tuning the parameters we were able to match the output of the model to several features of the recorded data, including symmetric amplitude modulation around zero and PSD plots as shown in Figures 7 and 8 (see further feature matching in Jones et al., 2009; Sherman et al., 2016), we arrived at the hypothesis that brief bouts of beta activity (“beta events”) non-time locked to alpha events could be generated by decreasing the mean delay between the proximal and distal drive to 0ms and increasing the strength of the distal drive relative to the proximal drive. This parameter set is also distributed with HNN (“AlphaandBeta.param”) and viewed in Figure 8A. With this mechanism, beta events emerged on cycles when the two stochastic drives hit the network simultaneously and when the distal drive was strong enough to break the upward flowing current and create a prominent ∼50ms downward deflection (see red box in Figure 8B). The stronger the distal drive the more prominent the beta activity (data not shown, see Sherman et al., 2016).

**Figure 8:**
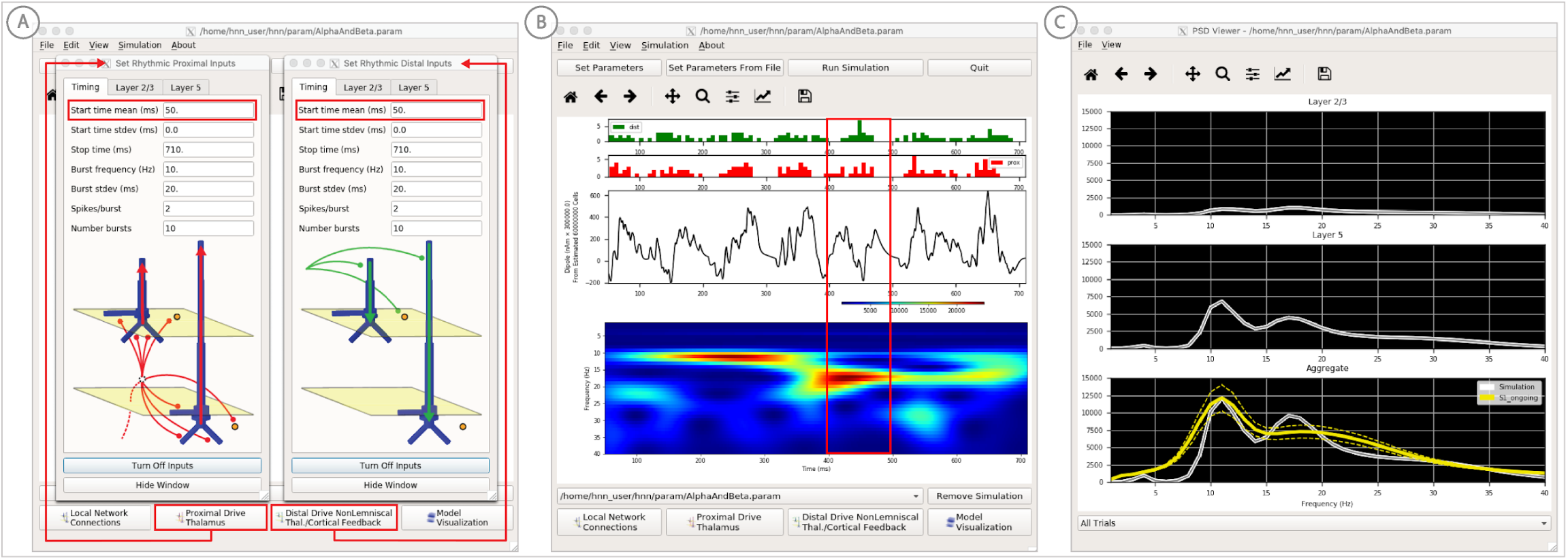
An example workflow for simulating transient alpha and beta frequency rhythm as in the spontaneous SI rhythms shown in Figure 6 (Jones et al 2009; Ziegler et al 2010; see Alpha and Beta Rhythms Tutorial text for details). **(A)** Here we are using the default HNN network configuration and not directly comparing the waveform to data, so begin with Step 3: activate the local network. In this example, a beta component emerged when the parameters of two ∼10Hz bursts to the local network through proximal and distal project pathways, as described in Figure 7, were adjusted so that on average they arrived to the network at the same time (see red boxes). This parameter set is provided in the “AlphaAndBeta.params” file. **(B)** Step 4: running the simulation with the “Run Simulation” button, shows that intermittent and transient alpha and beta rhythms emerge in the current dipole signal (middle dipole time trace; bottom time-frequency representation). Green and red histograms at the top display the defined distal and proximal burst drive patterns, respectively. Due to the stochastic nature of the bursts, on some cycles of the drive, the distal burst was simultaneous with the proximal burst and strong enough to push current flow down the dendrites to create a beta event (see red box). This model derived prediction reproduced several features of the data, including alpha and beta peaks in the corresponding PSD that were more closely matched to the recorded data (**C)**. Model predictions were subsequently validated with invasive recordings in mice and monkeys (Sherman et al 2016, see further discussion in text).

This beta event hypothesis was purely model derived and was based on matching several features of the SI mu rhythm between the model output and data (detailed in Jones et al., 2009; Sherman et al., 2016). One such feature was a direct comparison of the PSD between the model and data. This can be viewed in the HNN GUI though the “View PSD” pull down menu (see Figure 4, Step 5), where this data provided with HNN (“SI_ongoing.txt”) can be automatically compared the model output in the PSD window (Figure 8C).

The model derived predictions on mechanisms underlying alpha and beta where motivated by literature and further refined by tuning the parameters to match the output of the model with various features of the recorded data. While the mechanisms of the alpha rhythm described above were motivated by literature showing cortical alpha rhythms arise in part from alpha frequency drive from the thalamus and supported by animal studies (Hughes & Crunelli, 2005; for example, see Figure 2 in Bollimunta, Mo, Schroeder, & Ding, 2011), the beta event hypothesis was novel. The level of circuit detail in the model led to specific predictions on the laminar profile of synaptic activity occurring during beta events that could be directly tested with invasive recordings in animal models. One specific prediction was that the orientation of the current during the prominent ∼50ms deflection defining a beta event (red box, Figure 8B) was down the pyramidal neuron dendrites (e.g. into the cortex). This prediction, along with several others, were subsequently tested and validated with laminar recordings in both mice and monkeys, where it was also confirmed that features of beta events are conserved across species and recording modalities (Sherman et al., 2016; Shin et al., 2017).

#### Gamma rhythms

Gamma rhythms can encompass a wide band of frequencies from 30-150 Hz. Here, we will focus on the generation of so-called “low gamma” rhythms in the 30-80 Hz range. It has been well established through experiments and computational modeling that these rhythms can emerge in local spiking networks through excitatory and inhibitory cell interactions. The period of the low gamma oscillation is set by the time constant of decay of GABAA-mediated inhibitory currents (Buzsáki & Wang, 2012; Cardin et al., 2009; Vierling-Claassen, Cardin, Moore, & Jones, 2010), a mechanism that has been referred to as pyramidal-interneuron gamma (PING). In normal regimes, the decay time constant of GABA_A_-mediated synapses (∼25 ms) bounds oscillations to the low gamma frequency band (∼40 Hz). In general, PING rhythms are initiated by “excitation” to the excitatory (PN) cells, and this initial excitation causes PN spiking that, in turn, synaptically activates a spiking population of inhibitory (I) cells. These (I) cells then inhibit the PN cells, preventing further PN activity until the PN cells can overcome the effects of the inhibition ∼25 ms later. The pattern is repeated, creating a gamma frequency oscillation (∼40 Hz; 40 spikes/second).

We have applied HNN to determine if features in the current dipole signal could distinguish PING-mediated gamma from other possible mechanisms such as exogenous rhythmic drive (Lee & Jones, 2013). Here, we describe the process for generating gamma rhythms via the canonical PING mechanisms in HNN. We have not observed strong gamma rhythms in any of our prior studies. As such, while default parameters sets creating the example below are distributed with the software, no data are currently provided. We thus begin this tutorial with Step 3, “activating” the network using a slightly altered local network configuration, as described below.

**Step 3:** “Activate” the local network by loading in the parameter set defining the local network and initial input parameters “gamma_L5weak_L2weak.param”. In this example, the input was noisy excitatory synaptic drive to the pyramidal neurons. Additionally, all synaptic connections within the network are turned off (synaptic weight = 0), except for reciprocal connections between the excitatory (AMPA only) and inhibitory (GABA_A_ only) cells within the same layer. This is not biologically realistic, but was done for illustration purposes and to prevent pyramidal-to-pyramidal interactions from disrupting the gamma rhythm. To view the local network connections click the “local network” button in the Set Parameters dialog box. Figure 9B shows the corresponding dialog box where the values of adjustable parameters are displayed. Notice that the L2/3 and L5 cells are not connected to each other, the inhibitory conductance weights within layers are stronger than the excitatory conductances, and there are also strong inhibitory-to-inhibitory (i.e., basket-to-basket) connections. This strong autonomous inhibition will cause synchrony among the basket cells, and hence strong inhibition onto the PNs.

**Figure 9:**
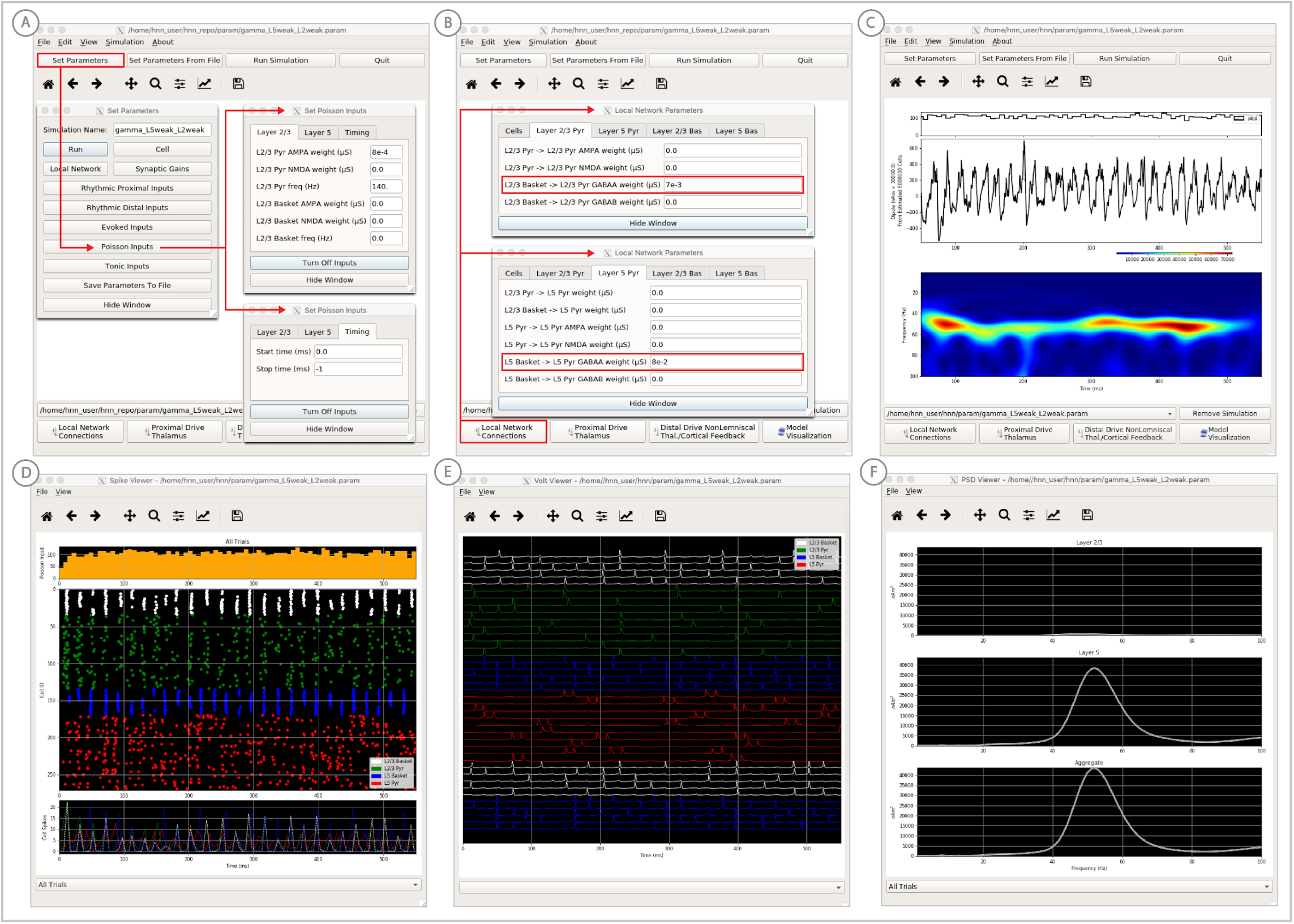
An example workflow for simulating pyramidal-interneuron gamma (PING) rhythms (Lee and Jones 2013; see Gamma Rhythms Tutorial text for details). **(A)** Here we are using the default HNN network configuration (with some parameter adjustments as shown in panel **B**) and we are not directly comparing the waveform to data, so begin with Step 3: activate the local network. Motivated by prior studies on PING mechanisms (see text), in this example PING rhythms were simulated by driving the pyramidal neuron somas with noisy excitatory synaptic input following a Poisson process. The parameters defining this noisy drive are viewed and adjusted through the Set Parameters button as shown, see text and Materials and Methods for parameter details. This parameter set for this example is provided in the “<u>gamma_L5weak_L2weak.param</u>” file. In this example, all synaptic connections within the network are turned off (synaptic weight = 0), except for reciprocal connections between the excitatory (AMPA only) and inhibitory (GABAA only) cells within the same layer. The local network connectivity can be viewed and adjusted through the Set Network Connection button or pull down menu, as shown in **(B)**. **(C)** Step 4: running the simulation with the “Run Simulation” button, shows that a ∼50Hz gamma rhythm is produced in the current dipole signal (middle dipole time trace; bottom time-frequency representation). The black histogram at the top displays the noisy excitatory drive to the network. **(D-F)** Step 5: additional network features, including cell spiking responses, somatic voltages, and layer specific power spectral density plots as shown can be visualized through the “View” pull down menu.

To reproduce the ∼40Hz gamma oscillation described by the PING mechanism above, we drove the pyramidal neuron somas in L2/3 and L5 with noisy excitatory AMPA synaptic input, distributed in time as a Poisson process with a rate of 140Hz. This noisy input can be viewed in the “Set Parameters” menu by clicking on the “Poisson Inputs” button (see Figure 9A). Setting the stop time of the Poisson drive to −1, under the Timing tab, keeps it active throughout the simulation duration.

**Step 4** The gamma rhythm shown in Figure 9C is reproduced by clicking the “Run Simulation” button at the top of the main HNN GUI. The top panel shows a histogram of Poisson distributed times of input to the pyramidal neurons, the middle panel the net current dipole across the entire network and the bottom the corresponding time frequency spectrogram showing strong gamma band activity. Additional network features, including spiking activity in each cell in the population (Figure 9D), somatic voltages (Figure 9E), and PSD plots for each layer and the entire network (Figure 9F), can also be visualized through the “View” pull down menu (**Step 5**). Notice the PING mechanisms described above in the spiking activity of the cells (Figure 9F), where in each layer the excitatory pyramidal neurons fire before the inhibitory basket cells. The line plots, which show spike counts over time, also demonstrate rhythmicity. The pyramidal neurons are firing periodically but with lower synchrony due to the Poisson drive (orange histogram at the top), which creates randomized spike times across the populations (once the inhibition sufficiently wears off). Notice also that the power in the gamma band is much smaller in Layers 2/3 than in Layer 5 (Figure 9F). This is reflective, in part, of the fact that the length of the L2/3 PNs is smaller than the L5 PNs, and hence the L2/3 cells produce smaller current dipole moments that can be masked by activity in Layer 5 (see Lee & Jones, 2013 for further discussion).

**Steps 6 and 7:** Local network and/or driving input parameters can be adjusted to explore alternate mechanisms of gamma generation and to develop and test hypotheses based on user defined data.

### ERP Model Optimization

To ease the process of narrowing in on parameter values representing a user’s hypothesized model, we have added a model optimization tool in HNN. Currently, this tool automatically estimates parameter values that minimize the error between model output and features of ERP waveforms from experiments. Parameter estimation is a computationally demanding task for any large-scale model. To reduce this complexity, we have leveraged insight of key parameters essential to ERP generation, along with a parameter sensitivity analysis, to create an optimization procedure that reduces the computational demand to a level that can be satisfied by a common multi-core laptop.

Two primary insights guided development of the optimization tool. First, exogenous proximal and distal driving inputs are the essential parameters to first tune to get an initial accurate representation of an ERP waveform. Thus, the model optimization is currently designed to estimate the parameters of these driving inputs defined by their synaptic connection strengths, and the Gaussian distribution of their timing (see dialog box in Figure 10B). In optimizing the parameters of the evoked response simulations to reproduce ERP data distributed with HNN (e.g. see ERP tutorial), we performed sensitivity analyses that estimated the relative contributions of each parameter to model uncertainty, where a low contribution indicated that a parameter could be fixed in the model and excluded from the estimation process to decrease compute time (see Supplementary Materials).

**Figure 10:**
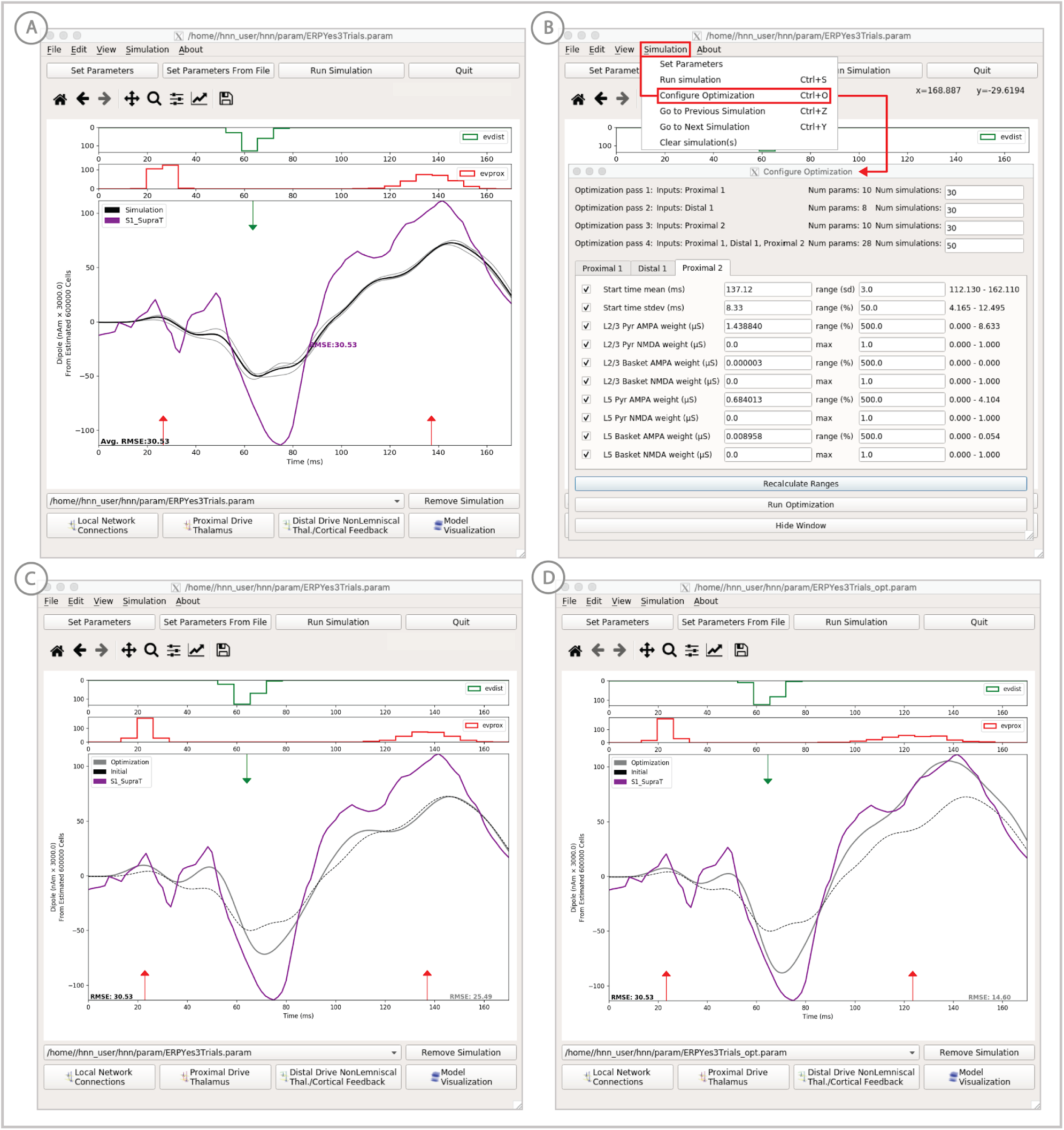
Example of the ERP parameter optimization procedure for a suprathreshold tactile evoked response. **(A)** Source localized SI data from a suprathreshold tactile evoked response (100% detection; purple) is shown overlaid with the corresponding HNN evoked response (black) using the threshold level evoked response parameter set detailed in Figure 4, as in initial parameter set. The RMSE between the data and the model is initially high at 30.53. **(B)** To improve the fit to the data, a serial procedure for optimizing the strengths of the proximal and distal drive input generating the evoked response can be run, by choosing the “Configure Optimization” option through the “Simulation” pull down menu. A dialog box allows users to choose and set a range over free optimization parameters, see text for details. **(C)** The GUI displays an intermediate fit after the first optimization step, specific to the first proximal drive. **(D)** The final fit is displayed once the optimization is complete. Here, the simulation from the optimized parameter set for the suprathreshold evoked response is shown in gray with an improved RMSE of 14.60 compared to 30.53 for the initial model. See Figure 10 Supplementary Figure 1 and 2 for further description of the optimization routine and a parameter sensitivity analysis.

Second, an intuitive insight that was confirmed by parameter sensitivity analysis is that the influence of each exogenous input on the simulated dipole varies over time, with the highest influence during and just after the time of the input (see Supplementary Material). We used this knowledge to create a stepwise optimization process, only estimating parameter values for one input at a time, where the objective of each optimization is to minimize a weighted root mean squared error (RMSE) measure between simulated and experimental data only during the relevant time window (see Materials and Methods). This stepwise estimation reduces the complexity of the optimization problem and saves time. Each step in the process searches for parameter estimates using the COBYLA optimization algorithm (Powell, 1994) (see Materials and Methods for detailed explanation of the stepwise optimization procedure).

### Example model optimization for the suprathreshold sensory evoked response data set

In this example, we describe an application of the model optimization tool for estimating parameters to simulate data representing the SI evoked response to a brief suprathreshold level tactile stimulation -- which is 100% detected (Figure 10A). This evoked response is similar to that shown in Figure 4, where the signal was elicited from a perceptual threshold level stimulation - at 50% detection. We start from the parameter file fitted to the 50% detection scenario, and use HNN’s model optimization feature to find parameter estimates that provide a better fit the suprathreshold-level experimental data. The data from this study is also included in the HNN distribution (“SI_SupraT.txt”).

**Steps 1-4:** Similar to steps 1-4 above, first load the supra-threshold experimental data file “S1_SupraT.txt” via the “Load data file” menu option and the example starting parameters to activate the network provided in the parameter file “ERPYes100Trials.param” via the “Load parameter file” menu option. Note that in this example, the network is also “activated” by a sequence of three exogenous inputs defined in the parameter file. The parameters for these inputs serve as a baseline for model optimization. The supplied parameter file (used above) runs 100 trials by default for each simulation. For model optimization, this can be reduced to 3 trials. Click on the “Set Parameters” button, then the “Run” button, and replace 100 trials with 3. In the previous Set Parameters dialog box change the simulation name to “ERPYes3Trials” to reflect this change (Figure 4C). By clicking the “Run Simulation” button the evoked response using this initial parameter set as in Figure 10A will be displayed. As described above, in practice with user defined data, users should apply their own hypotheses related to the number, timing and synaptic input strengths of the exogenous inputs that activate the network to obtain an initial representation of the recorded waveform before beginning the parameter estimation process.

**Step 5**: Before running the optimization, rename the simulation to “ERPYes3Trials_opt” in the Set Parameters dialog box as described above, so that the parameter results of the optimization will be saved in a new file.

**Step 6:** In the Simulation pull down menu, choose the “Configure Optimization” option. This option is only selectable once data and parameter files have been loaded. A new dialog box pre-populated with values from the parameter file will appear, as shown in Figure 10B. All parameters describing the timing and strength of defined exogenous inputs will be available for optimization. Users can generate their own evoked response parameter files with as many exogenous inputs as desired and they will be automatically populated into the “Configure Optimization” dialog box.

Select which parameters to treat as free variables for optimization; parameters that will be fixed in the optimization process are grayed out. By default, all parameters are selected, but it may be desirable to limit the number of free parameters to only the most influential set based on a parameter sensitivity analysis. Fixing non-influential parameters will decrease the complexity of an optimization step, and increase the likelihood of the optimization algorithm converging on parameter estimates after a relatively low number of simulations. Results of a sensitivity analysis using Uncertainpy (Tennøe, Halnes, & Einevoll, 2018) on this example data are provided in Supplementary Table 1 and may help guide model optimization for similar data. Sensitivity analysis is not yet included in HNN (see Future Directions).

The number of simulations per optimization step is configurable in the top section of the “Configure Optimization” dialog box (Figure 10B). The default values shown in Figure 10B were based on results from our studies where the fit obtained was significantly improved from a single optimization. This value can be decreased as the number of free parameters is reduced.

The parameter ranges defining the bound constraints given to the optimization algorithm are shown in the right-most column of the dialog box in Figure 10B. The displayed range is calculated as plus or minus a specified number of standard deviations for input start time or plus or minus a percentage of the initial value for all other parameters. If a parameter has an initial value of 0, its range is defined by a user-specified maximum value rather than percentage. The “Recalculate Ranges” button will display updated values.

**Step 7**: Click the “Run Optimization” button to start the stepwise optimization process.After each input has been optimized in sequence and a final optimization pass over all parameters has completed, the final optimized fit will be shown in gray in the main HNN window along with the lowest obtained RMSE (Figure 10C/D).

**Step 8 (optional)**: To perform a second optimization using the results of the first procedure as a starting point, select the optimized simulation parameter set drop-down menu. This will update the values in the Configure Optimization dialog box and pressing Run Optimization will start a new optimization process. For this example, the RMSE improved from 14.60 (Figure 10D) after the first optimization to 10.79 after a second round (data not shown).

## Discussion

The Human Neocortical Neurosolver (HNN, https://hnn.brown.edu) is a neural modeling software tool developed to help researchers and clinicians interpret the neural origin of their human EEG or MEG data. HNN’s interactive GUI is designed for users with no formal computational neural modeling or software development experience to be able to develop and test hypotheses on the cellular- and circuit-level generators of their human data. Based on prior applications of HNN’s underlying template neural model on these signals (Jones et al., 2009, 2007; Khan et al., 2015; Lee & Jones, 2013; Sherman et al., 2016; Sliva et al., 2018; Ziegler et al., 2010), the tutorials and the example workflow focus on studying the neural origin of ERPs and low-frequency oscillations from a single brain region. The template network model contains features of a canonical neocortical circuit, with layer-specific thalamocortical and cortico-cortical drive, where the net primary current dipoles are simulated from the intracellular current across the network of pyramidal neurons. HNN enables visualization and direct comparison of the primary current dipole produced by the network to source-localized data in units of nAm, under various parameter manipulations. This comparison, along with simultaneous visualization of microcircuit activity, including cell spiking and somatic voltage responses, guides interpretation of the cellular- and circuit-level origin of EEG/MEG data.

HNN was created based on the biophysical origin of EEG/MEG primary currents to be a hypothesis development and testing tool, where specific predictions on the microcircuit-level underpinnings of recorded data can be produced. The circuit-level predictions can guide further validation with invasive recordings or with other imaging modalities (e.g., spectroscopy or tractography, see Khan et al., 2015). As one specific example, HNN led to a novel prediction on the origin of transient neocortical beta oscillations, and the prediction was later tested and supported by laminar recordings in mice and monkeys (Sherman et al., 2016). In turn, established cellular- or circuit-level details known to contribute to healthy brain dynamics and/or disease states can be adapted into HNN to predict corresponding signatures in macroscale signals.

HNN is particularly timely given the rapidly expanding wealth of genetic insights and phenotype data in animal model systems. As disease-specific genetic mutations and corresponding cellular/circuit outcomes in mouse models are identified, they can be implemented in HNN, and their impact on EEG/MEG measured brain dynamics, ranging from ongoing state properties (e.g., alpha oscillations) to sensory-evoked responses, can be simulated. The outputs from HNN would then provide specific and principled predictions to be compared against real EEG/MEG data obtained in the relevant population, leading to valid bi-directional inference. Overall, the scalability of HNN provides an unprecedented framework for translational neuroscience research.

### Comparison to other EEG/MEG modeling software

Although other models and software packages aimed at providing researchers tools to simulate macroscale EEG/MEG signals are available (e.g., https://thevirtualbrain.org; https://www.fil.ion.ucl.ac.uk/spm/ (Barrès, Simons, & Arbib, 2013; Hagen, Næss, Ness, & Einevoll, 2018; Kiebel, Garrido, Moran, & Friston, 2008; Sanz Leon et al., 2013). HNN’s model, goals, and capabilities are unique in this realm. To our knowledge, HNN is the only software able to simulate the primary electrical currents underlying EEG/MEG signals from the intracellular dendritic current flow in multi-compartmental pyramidal neurons embedded in a detailed model of layered cortical circuitry that contains individual spiking neurons and layer-specific drive from thalamocortical and cortico-cortical networks. This construction was specifically designed for interpreting microscale cellular- and circuit-level activity from single regions of interest.

Other models typically rely on reduced representations of neural activity, including neural mass representations and/or mean field approximations (Breakspear, Williams, & Stam, 2004; Jansen & Rit, 1995; Jirsa & Haken, 1996; Kiebel et al., 2008; Sanz Leon et al., 2013; Woolrich & Stephan, 2013). Such simplifications may be necessary to ensure mathematical or computational tractability of models that address whole brain activity or interactions between multiple areas (Breakspear, 2017), but that tractability comes at the cost of suppressing or eliminating the ability to evaluate the roles of cellular-level details of individual spiking units or particular network architectures, or to perform one-to-one comparisons between model and data.

In contrast to the more simplified models mentioned above, the biophysical detail included in HNN’s model enables direct comparison between model output and source-localized data in the same units of measure (nAm), facilitating GUI-driven hypothesis development and testing. Additionally, unlike other software, HNN’s GUI, workflow, and tutorials are uniquely designed to train users to stimulate ERPs and oscillations based on layer-specific exogenous drive to the network, as validated in prior studies (see Introduction). Although HNN’s level of detail can be prohibitive for studying more global network interactions where the computations are more tractable with reduced neural representations, the level of detail built into HNN is necessary to make predictions on local microscale cellular and circuit elements contributing to EEG/MEG that can guide targeted testing and direct connection to studies in animal models.

### Limitations and Future directions

One of the greatest challenges in computational neural modeling is deciding the appropriate scale of model to use to answer the question at hand. There is always a tradeoff between model complexity and computational efficiency, ease of use, and interpretability. As discussed above, this tradeoff underlies different scales of modeling in various EEG/MEG modeling software. HNN’s model was chosen to be minimally sufficient to accurately account for the biophysical origin of the primary currents that underlie EEG/MEG signals; namely, the net intracellular current flow in the apical dendrites of pyramidal neurons that span across the cortical layers and receive layer-specific synaptic input from other brain areas. While HNN’s model is a reduction of the full complexity of neocortical circuits, it has been successful in interpreting the origin of extracranially measured macro-scale EEG/MEG signals that likely rely on canonical features of neocortical circuitry and do not provide access to finer details of the underlying structure. That said, any conclusions made with HNN are based on the underlying model assumptions that are important for users to understand. These assumptions are outlined in detail in the methods, in our prior publications, and on our website.

Parameter optimization is a computationally challenging problem in any large-scale model. The process for parameter tuning to study ERPs and oscillations in HNN’s underlying model is detailed above. Based on our prior studies and sensitivity analyses (see Supplementary Materials), we have identified that the timing and strength of the layer specific exogenous drive to the local network is critical in defining the timing and peaks of sensory evoked responses. As such, HNN currently includes a tool to optimize these parameters based on reducing the error between simulated evoked response waveforms and recorded data. Due to the non-stationary nature of spontaneous brain rhythms (e.g. Figure 6) error reduction based on matching waveform features is not as straightforward, and other signal features may be necessary to consider for optimization (e.g. PSD peak amplitudes, see Figure 8 and Jones et al., 2009). Future expansions of HNN will include the ability to optimize over other user defined parameters, and to minimize errors between model output and various features of recorded data, with an estimate of the sensitivity of various parameters to these features. Given enough compute power, large parameter sweeps could be implemented in HNN to generate families of models could that could then be template matched to given waveforms via machine learning algorithms, as an alternative means for circuit interpretation without interactive hypothesis development and testing. At present, HNN can be run on high performance computers through the Neuroscience Gateway Portal (www.nsgportal.org) and Amazon Web Services (https://aws.amazon.com), see also Dissemination in Materials and Methods.

Currently, all conclusions made in HNN are derived from the template neocortical column model provided. Another important step in expanding HNN’s utility will be to enable users to define their own cells and circuits to use within the HNN framework. While the HNN code is open source and adaptable for advanced users, it is difficult for those without expertise in computational neural modeling in Neuron/Python to expand. Therefore, work is in progress to convert HNN’s underlying neural model to the NetPyNe simulation language (www.netpyne.org) (Dura-Bernal et al., 2019). NetPyNe is a neural modeling platform enabling flexible cell and network development. This conversion will also facilitate the ability to expand HNN to the study of activity from and between multiple cortical areas and the thalamus.

HNN is designed to simulate source-localized current dipole signals produced by neurons. Source localization is currently viewed as an independent process. The output from any source localization algorithm can be compared to HNN’s simulated output. In future expansions of HNN, we plan to integrate HNN’s “bottom up” simulations, with “top down” source localization estimates using minimum-norm-estimate (MNE) software (www.martinos.org/mne) (Gramfort et al., 2013, 2014), providing an all-in-one software tool for source localization and circuit-based interpretation. In doing so, parameter estimation in each software package may benefit from direct knowledge and constraints from the other. Additionally, HNN’s utility will be expanded to include estimation of forward fields through the brain to simulate and visualize local field potential, current-source density, and sensor-level signals, facilitating comparison to these recording modalities.

We have shown that HNN can be a useful tool to interpret the impact of noninvasive brain stimulation (NIBS) on EEG-measured circuit dynamics (Figure 5, Sliva et al., 2018). HNN was used to test specific hypotheses on tACS-induced modulation of synaptic dynamics by accounting for EEG signal differences in pre-tACS compared to post-tACS periods. A useful expansion of HNN will be to include simulations of the fields induced in the brain by NIBS (e.g., with finite-element-estimates (Windhoff, Opitz, & Thielscher, 2013)) and to directly couple these fields to the modeled neurons. This integration would facilitate studying the effects of NIBS on real-time EEG signals and could lead to improved NIBS paradigms.

In total, HNN’s present distribution and planned expansions are aimed at providing a one-of-a-kind, user-friendly software tool for translational neuroscience research that is accessible to a wide scientific and clinical community.

## Materials and Methods

### Dissemination

HNN is distributed online at https://hnn.brown.edu. The menu bar at the top of the HNN homepage (Figure 11) links to installation instructions, documentation, tutorials, troubleshooting information, and a user forum. HNN can be installed locally on Linux, Windows, and macOS operating systems, and it can be run as well online through Amazon Web Services (AWS) or the Neuroscience Gateway Portal (NGP). Since HNN is an open-source project, the code for our software, as well as the local installation instructions, are hosted on GitHub (see https://github.com/jonescompneurolab/hnn).

**Figure 11:**
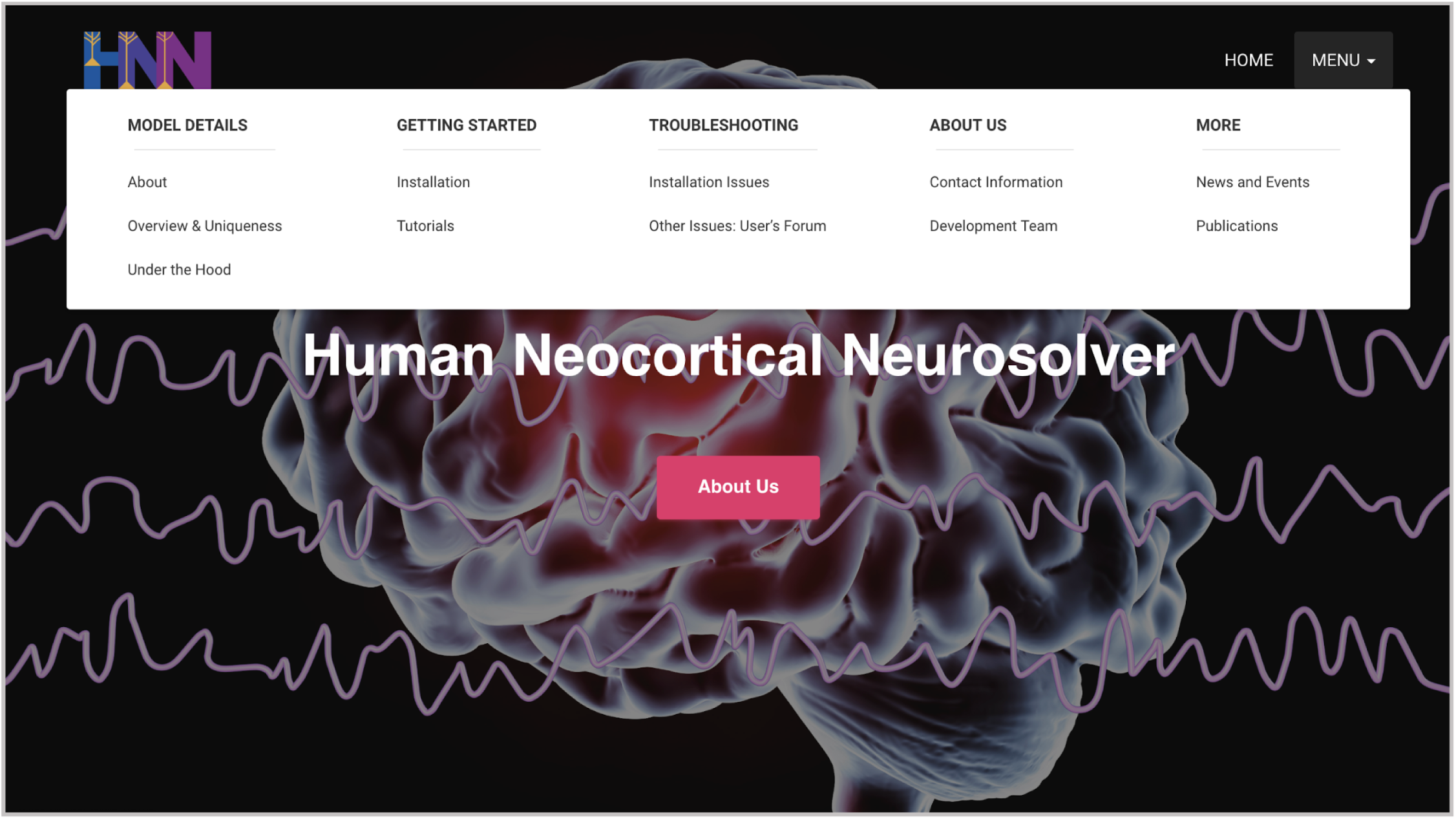
The homepage of the HNN website http://hnn.brown.edu and menu items containing installation instructions, documentation, tutorials, and troubleshooting information.

### Template Model Construction

#### Overview

The template neocortical model provided with HNN is based on prior publications using the model without a graphical user interface, as described in Jones et al., 2009, and available on ModelDB (https://senselab.med.yale.edu/modeldb). All current parameter files included in the software, and described in the tutorials above, are based on this model, except for the gamma tutorial whose parameter file has local network modifications as described above.

HNN’s underlying neocortical model is simulated using the NEURON simulation environment with the Python interpreter. HNN’s model is simulated across multiple cores in parallel using the message-passing interface (MPI). HNN’s Run Parameters dialog box can be accessed through the GUI and provides access to commonly used simulation parameters, including integration time-step (dt), simulation duration (milliseconds), number of trials, neuronal firing threshold (mV), and number of cores over which to parallelize the model.

The model represents a canonical neocortical circuit. It contains multi-compartment pyramidal neurons (PN) in supragranular and infragranular layers (layers 2/3 and 5, respectively), whose apical dendrites are spatially aligned and span the cortical layers. In both layers, the PNs have two basal, one oblique, and one apical dendrite branch, and the layer 2/3 PNs have shorter apical dendrites than layer 5 PNs.

The PNs are synaptically coupled to each other and to a subset of inhibitory neurons in each layer, and are included in the model in a 3/1 PN-to-interneuron ratio, with a scalable number of PNs. The inhibitory neurons are simulated with single compartments representing fast spiking basket cells, and are shown in yellow in (Figure 2). Note that the granular layer is not explicitly included in the template circuit. This design choice was based on the fact that macroscale current dipoles are dominated by PN activity in supragranular, and infragranular layers, due to their alignment (see Calculation of Primary Electrical Current). Thalamic input to granular layers is presumed to propagate directly to basal and oblique dendrites of PN in layer 2/3 and 5.

### Detailed Neuronal Morphology and Physiology

#### Morphology

The morphology of the PN in each layer (see Figure 3, and Table 1) were adapted from the morphology reduction procedure in (Bush & Sejnowski, 1993), which was based on conserving the axial conductance between compartments in the cells. The axial conductance is the basis of the primary current dipole calculation (see below).

**Table 1.**
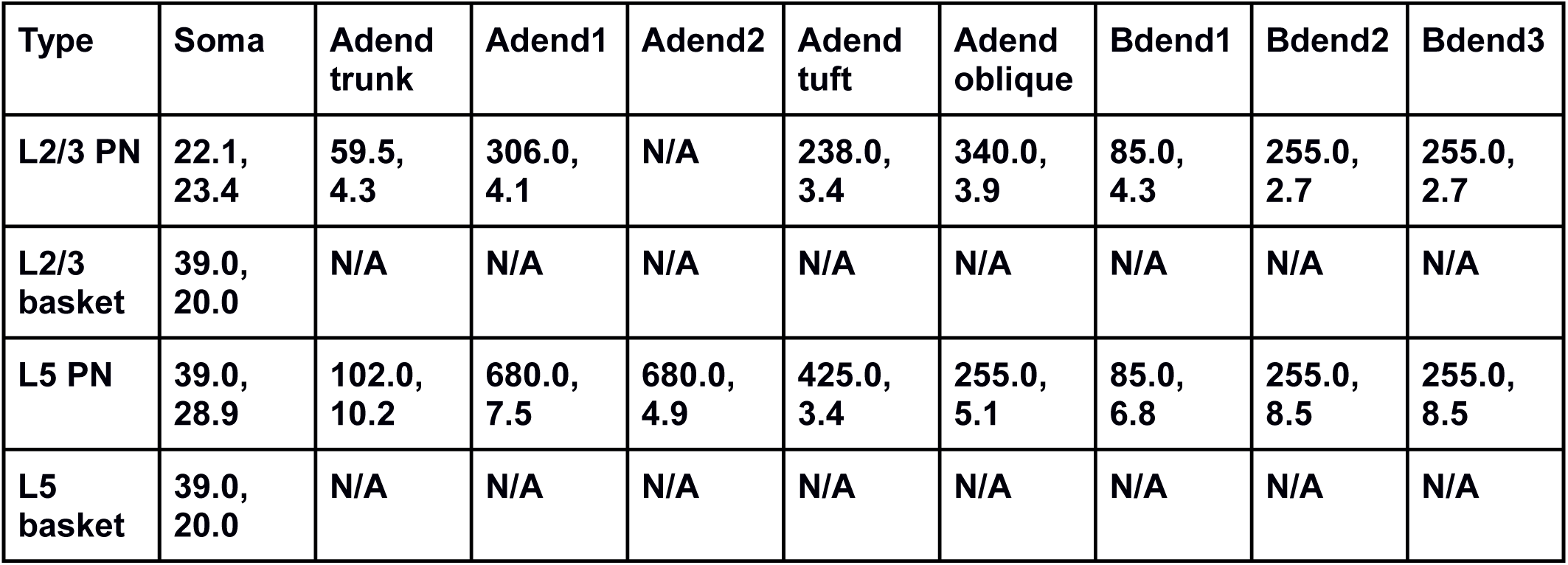
Length (*μm*), diameter (*μm*) of compartments in each modeled neuron type. Adend (Bdend) represent apical (basal) dendrite. Note the following connectivity for compartments of PNs. The non-oblique Adends of PNs are connected vertically along the Z axis (cortical layer axis from supra- to infragranular layers) from soma → Adend trunk → Adend1 → Adend2 → Adend tuft. The smaller L2/3 PNs do not have an Adend2 compartment. PN Adend oblique are connected to the soma and perpendicular to the Z axis. Bdend1 connects to the soma along the Z axis, and Bdend2 and Bdend3 branch from Bdend1 at a 45 degree angle from the Z axis. L2/3 and L5 basket interneurons have a single somatic compartment. N/A indicates non-applicable, since that specific compartment not present in the neuron type. Geometry illustrated in Figure 3 above.

| Type | Soma | Adend trunk | Adend1 | Adend2 | Adend tuft | Adend oblique | Bdend1 | Bdend2 | Bdend3 |
| --- | --- | --- | --- | --- | --- | --- | --- | --- | --- |
| L2/3 PN | 22.1, 23.4 | 59.5, 4.3 | 306.0, 4.1 | N/A | 238.0, 3.4 | 340.0, 3.9 | 85.0, 4.3 | 255.0, 2.7 | 255.0, 2.7 |
| L2/3 basket | 39.0, 20.0 | N/A | N/A | N/A | N/A | N/A | N/A | N/A | N/A |
| L5 PN | 39.0, 28.9 | 102.0, 10.2 | 680.0, 7.5 | 680.0, 4.9 | 425.0, 3.4 | 255.0, 5.1 | 85.0, 6.8 | 255.0, 8.5 | 255.0, 8.5 |
| L5 basket | 39.0, 20.0 | N/A | N/A | N/A | N/A | N/A | N/A | N/A | N/A |

*Layer 2/3*

- PN: 8 compartments including 4 apical dendrites, 3 basal dendrites, 1 soma
- Inhibitory basket neurons: single compartment (soma)

*Layer 5*

- PN: 9 compartments including 5 apical dendrites, 3 basal dendrites, 1 soma.
- As shown below, L5 PNs have longer dendrites than L2/3 PNs. L5 PN somas are based in L5 with long apical dendrites reaching into L2/3.
- Inhibitory Basket neurons: single compartment (soma).
- L2/3 and L5 basket interneurons are identical but their synaptic parameters and local circuit connectivity differs.

### Physiology

Membrane voltages in each simulated compartment were calculated using the standard Hodgkin-Huxley parallel conductance equations, and current flow between compartments follows from cable theory as accounted for in NEURON.

Active ionic currents were included in both the somatic and dendritic compartment of the cells. The parameters regulating these currents were tuned to replicate known *in vitro* firing patterns in response to somatic current injection. The L5 PNs produced bursts of action potentials after sufficient depolarization and the L2/3 PNs produced adapting spike trains.

The following table displays the ion channels and mechanisms in each cell type in the model (**X)** indicates the presence of the channel/mechanism in the cell type, see online code for full equations.

| Cell Type (rows)<br>Channel/<br>mechanism Type<br>(columns) | Na<br>(fast) | K<br>(fast) | Km | KCa | Ca (L-type) | Ca (T-type) | Ca<br>decay | HCN | leak | dipole |
| --- | --- | --- | --- | --- | --- | --- | --- | --- | --- | --- |
| Basket | X | X |  |  |  |  |  |  | X |  |
| L2/3 Pyramidal | X | X | X |  |  |  |  |  | X | X |
| L5 Pyramidal | X | X | X | X | X | X | X | X | X | X |

In the table above, Na (fast) / K (fast) are the fast sodium and potassium channels responsible for generating action potentials. Km is the muscarine sensitive potassium channel, with a relatively slow time-constant and KCa is the calcium-dependent potassium channel, which contributes to hyperpolarization after calcium influx into the cell. The L- and T-type calcium (Ca) channels represent the high-threshold and low-threshold activated calcium channels which together with the hyperpolarization-activated cyclic nucleotide gated channel (HCN) contribute to bursting. Ca decay represents the calcium extrusion pump, which causes intracellular calcium to decay towards a baseline level. Leak represents the passive channel, with constant conductance. Dipole represents the mechanism that takes into account the primary axial current flow within pyramidal neuron dendrites, responsible for the generation of simulated signals comparable to MEG/EEG recordings. For more details see Jones et al., 2009.

### Local Network Connections

HNN’s default template neocortical model includes neurons arranged in three dimensions. The *XY* plane is used to array cells on a regular grid while the Z-axis specifies cortical layer. HNN’s default model contains a regular 10 x 10 grid (arbitrary units) of pyramidal neurons in layer 2/3 and layer 5 for a total of 200 pyramidal neurons, with interneurons interleaved regularly in a 3-1 ratio (see Figure 3D).

Synaptic dynamics were modeled with bi-exponential functions. The rise and decay time constants and reversal potentials were based on experiments and the original neocortical model in Jones et al., 2009, and are generally as follows: AMPA (0.5 ms, 1.0 ms, 0 mV); NMDA (1.0 ms, 20.0 ms, 0 mV); GABAA (0.5 ms, 5.0 ms, −80 mV), GABAB (1.0 ms, 20.0 ms, −80 mV). Within a cortical layer there is recurrent connectivity between neurons of a given type (PN to PN, interneuron to interneuron), PN to interneuron connectivity, and synaptic inhibition from interneurons onto PNs. The following synaptic connections are present across cortical layers: layer 2/3 PNs to layer 5 PNs, layer 2/3 interneurons to layer 5 PNs, layer 2/3 PNs to layer 5 interneurons.

There is all-to-all connectivity between any two populations of synaptically-coupled neurons. Synaptic *weights* between the neurons are scaled inversely by the distance in the XY plane (arbitrary units) between the neurons (*d*) using exponential fall-off following *e*^−*d*^2^/*λ*^2^^, and space constant *λ*, which depends on pre- and post-synaptic type (Table 2 below).

**Table 2.**
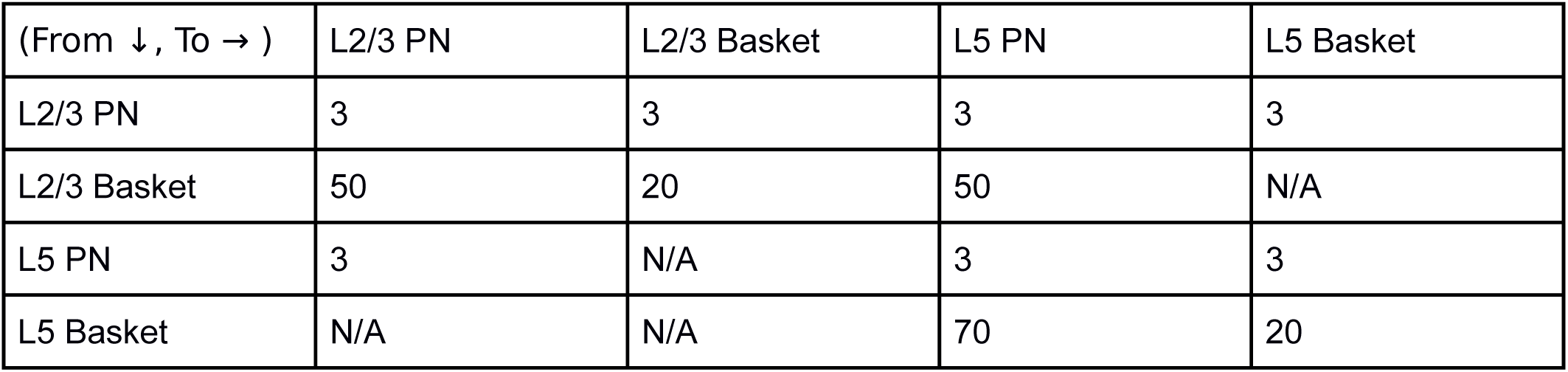
Space constant (arbitrary units) for synaptic connection strengths and delays between different populations of neurons (rows are pre-synaptic type, columns are post-synaptic type).

| (From ↓, To → ) | L2/3 PN | L2/3 Basket | L5 PN | L5 Basket |
| --- | --- | --- | --- | --- |
| L2/3 PN | 3 | 3 | 3 | 3 |
| L2/3 Basket | 50 | 20 | 50 | N/A |
| L5 PN | 3 | N/A | 3 | 3 |
| L5 Basket | N/A | N/A | 70 | 20 |

The synaptic *delays* are scaled in proportion to the XY plane distance (*d*) between the neurons following 1/*e*^−*d*^2^/*λ*^2^^, to account for the larger propagation distance (note that the *λ* value is determined using values in Table 2). With increasing *d* between neurons, the synaptic weights decay, while the synaptic delays increase. The connectivity details are based on known neocortical anatomy and local circuit wiring patterns, as derived from the literature. Further details on connectivity are available on HNN’s website and prior publications.

### Exogenous Driving Inputs

At rest, the default model does not generate activity. HNN provides several ways to activate the local cortical column with layer specific excitatory synaptic input representing thalamo-cortical, and/or cortical-cortical and noisy/tonic drive. The user defines the choice of driving input to the network, based on their simulation experiment, as described in Results.

Exogenous driving networks are not explicitly modeled, rather the user defines trains or bursts of action potentials representing these inputs that excite the local network via AMPA or NMDA synaptic connections to distinct layers and cellular compartments. These inputs are referred to as proximal and distal drive based on the PN dendritic contact location. Proximal inputs contact basal and oblique dendrites of PN and somas of the inhibitory neurons in L2/3 and L5, and distal inputs contact distal dendrites of the PN in L2/3 and L5 and somas of the inhibitory neurons in L2/3 only, as shown in Figure 3.

The trains of action potentials, or tonic/noisy input, that the user defines are created in specific dialog boxes in the GUI and represent either Evoked, Rhythmic, Tonic, or Poisson Inputs, as motivated by our prior studies and tutorials described in Results.

#### Evoked Input

Evoked inputs are trains of synaptic inputs to the local network during a sensory stimulus that creates an event related potential (ERP). Parameter choices for defining these inputs are shown in Figure 4A. The following parameter values are used to define each proximal or distal evoked input:

- Start time mean (ms) - average start time
- Start time stdev (ms) - standard deviation of start time
- Number spikes - number of inputs provided to each synapse
- L2/3 Pyr weight AMPA/NMDA (*μS*) - weight of AMPA/NMDA synaptic inputs to layer 2/3 pyramidal neurons
- L2/3 Basket weight AMPA/NMDA (*μS*) - weight of AMPA/NMDA synaptic inputs to layer /32 basket cells
- L5 Pyr weight AMPA/NMDA (*μS*) - weight of AMPA/NMDA synaptic inputs to layer 5 pyramidal neurons
- L5 Basket weight AMPA/NMDA (*μS*) - weight of AMPA/NMDA synaptic inputs to layer 5 basket cells (only used for *proximal* inputs)

Each evoked input also has a “Synchronous Inputs” option, indicating whether for a specific evoked proximal/distal input each neuron receives the input at the same time, or if instead each neuron receives the evoked input events independently drawn from the same distribution. Increment input (ms) indicates whether to increment the Start time of all evoked inputs on each trial. In the studies described above, the evoked input strengths are suprathreshold generating action potentials in the local network.

#### Rhythmic Input

Rhythmic Inputs are typically bursts of action potentials that drive the local network rhythmically. Parameter choices for defining these inputs are shown in Figure 7A. Each rhythmic input is defined as a series of “population bursts”, consisting of a set number of “burst units” which drive post-synaptic conductances in the local network with a set frequency and mean delay between proximal and distal projections. Rhythmic proximal and distal inputs target different cortical layers, as described above. HNN allows setting proximal and distal rhythmic synaptic input start/stop times and frequencies using the following specification:

- Start time mean (ms) - specifies the average start time for rhythmic inputs
- Start time stdev (ms) - specifies the standard deviation of start times for rhythmic inputs
- Stop time (ms) - specifies when the rhythmic inputs should be turned off
- Burst frequency (Hz) - average frequency of bursts
- Burst stdev (ms) - standard deviation of input events
- Spikes/burst - provides *n* synaptic events at each selected time
- Number bursts - number of times the full Burst sequence is repeated (each repeat adds variability and more inputs)

In addition, HNN’s Rhythmic Input dialog box allows setting the weights of the rhythmic synaptic inputs (units of conductance) to individual neuron types in layers 2/3 and 5, and adding synaptic delays (ms) before the neurons receive the synaptic inputs. In the studies described above, rhythmic inputs are set to sub-threshold synaptic strengths, and therefore do not lead to neuronal action potentials.

#### Tonic/Noisy Input

Tonic inputs are modeled as somatic current clamps with a fixed current amplitude (nA). These clamps can be used to adjust the resting membrane potential of a neuron, and bring it closer (with positive amplitude injection) or further from firing threshold (with a negative amplitude injection). Parameter choices for defining these inputs are shown in Figure 9A and include setting the current clamp amplitude, and start/stop time for each modeled neuron type separately.

Noisy Inputs are trains of action potentials that follow a Poisson Process and create excitatory AMPA or NMDA synaptic inputs to the somata of all neurons of a given type. Parameter choices for defining these inputs are shown in Figure 9A and include, setting the average frequency of the Poisson drive, synaptic strength to somatic AMPA or NMDA synapses, and start/stop times of all Poisson inputs.

### Calculation of Primary Electrical Current (Net Current Dipole)

Axial current flow between any two neighboring model compartments i,j is defined as i_axial_ = (v_i_ - v_j_) / r_axial_, where v_i_, v_j_, and r_axial_ are the voltages in compartment i, j, and the resistance between the compartments, respectively. In order to convert this axial current into a dipole signal, we apply a length scaling where the axial current is scaled by the inter-compartment distance along the vertical axis. The length scaling means that for the longer apical dendrites of layer 5 pyramidal neurons, the contribution will be larger than from the shorter layer 2/3 pyramidal neuron apical dendrites. Note that the orientation of the dendrites relative to the vertical axis also influences the contribution to the dipole signal. For example, the horizontally-oriented oblique dendrites which do not have any vertical length component, do not contribute to the dipole signal, whereas for basal dendrites oriented at 45 degrees from the vertical axis, the scaling is -√2/2 (note the negative sign is because these dendrites are pointing downward). The contribution from all neighboring compartments within a neuron is integrated and then added to a value across the set of all pyramidal neurons. As a result of the multiplication between axial current and length, the model dipole output signal has the same units of measure as the experimental data in units of nanoAmpere-meters: nAm (Okada et al., 1997).

### ERP Optimization Tools

HNN includes a method to optimize ERP simulations. The optimization procedure was uniquely designed to minimize the RMSE between model output and ERP waveforms in a stepwise manner that decreases parameter exploration and saves compute time. This procedure takes advantage of the assumption that the exogenous proximal and distal driving inputs are essential parameters to tune to get an accurate representation of an ERP waveform. Additionally, it applies the knowledge that, with probabilistic certainty, features of the dipole waveform at a particular point in time cannot be influenced by an exogenous driving input that begins after that point in time.

Since exogenous inputs are modeled as Gaussian processes, the likelihood of occurrence can be modeled by a probability distribution function (PDF) normally distributed with a given mean and standard deviation. Figure 10 Supplemental Figure 1A shows the PDFs of the inputs for the suprathreshold example described in the results Figure 10. An input’s contribution to the ERP will begin when there is a non-zero probability of occurrence and persist for a duration commensurate with the input’s cumulative distribution function (CDF), shown in Figure 10 Supplemental Figure 1B. This clearly illustrates that from 20-50 ms, the input labeled “Proximal 1” is the unique contributor to the waveform. After 50 ms, effects from Distal 1 begin, thus adding new parameters that contribute to the waveform fit and reduce the relative contribution of Proximal 1 (from full to partial). It follows that each successive driving input will have a time window where it is most likely to have a unique and dominant effect. As such, our approach to model optimization is to divide the process into smaller steps where only a single input’s parameters are estimated before proceeding to optimize the next input.

To implement this procedure, we developed a new goodness of fit measure that amplifies the importance of maximizing the fit at points of unique contribution (e.g. 20-50 ms for Proximal 1, Figure 10 Supplemental Figure 1C) and diminished the importance of fitting to later points where other inputs contribute more to the fit. We began with standard root mean squared error (RMSE)

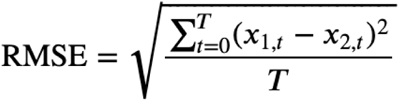

where *t* is the current simulation time, from 0 to simulation completion (*T*), and *x_1,t_* is the simulated dipole at *t,* and *x_2,t_* is the experimental data point. Then we adapted RMSE to include weight functions specific for input *k* at time *t*,

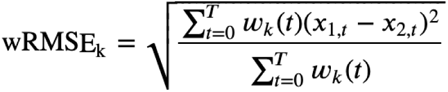

where an assignment of *w_k_(t)* = 1 for all *t* would be equivalent to RMSE.

For each input *k*, we first defined a weight distribution function, *w_k_(t)*, as the Unique Contribution Index (UCI), which starts from the CDF of input *k* and simply subtracts the CDF of subsequent inputs, with a lower bound of 0 (Figure 10 Supplemental Figure 1C). Equivalently,

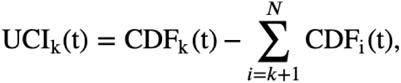

where *N* is the number of exogenous driving inputs in the simulation. Figure 10 Supplemental Figure 1C shows that Proximal 1’s influence is unique up to 50ms, Distal 1 has a dominant, but not unique contribution near 70 ms, and Proximal 2 is dominant after ∼100 ms. When the UCI is applied as a weighting function in the wRMSE equation above, we observed that some optimization steps would negatively impact the fit in regions after the peak in UCI, where the errors had been down-weighted, requiring subsequent optimization steps to attempt to “correct” the fit. Our solution was to instead define the weight function using the Extended Contribution Index (ECI), which includes a term that delays the weight function’s return to 0, extending the window of data points that have an impact on wRMSE further into the simulation. This achieves a balance between optimal parameter estimates for the current step and providing a good starting point for following optimization steps. ECI is defined by

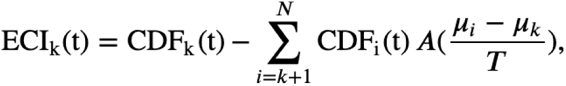

where μ_i_ and μ_k_ are the mean start times of the next input and the current input, respectively. Simulation length is represented by *T* and *A* is an empirically derived constant. We arrived at a value of 1.6 for *A* as a factor that appropriately minimized the contribution of inputs proportional to the delay between their onset and the *k*^th^ input currently being optimized. The effect of the ECI’s decay term can be seen in Figure 10 Supplemental Figure 1D where the ECI for Proximal 1 extends further than the corresponding UCI, and the ECI of Distal 1 remains significant through the end of the simulation. Since points where ECI_k,t_ approximately equal 0 will have a negligible impact on wRMSE_k_, we define a threshold of 0.01 where wRMSE_k_ is calculated for the window starting when ECI_k,t_ rises above 0.01 and ending when ECI_k,t_ drops below 0.01. For the first exogenous driving input, it is likely that the window will end before the completion of the simulation. In that first step, simulations can be stopped early, reducing the time required for simulating each candidate parameter set in that step.

The final step in our model optimization process is to vary all free parameters from all inputs using regular RMSE to measure goodness of fit. Like each previous step, the number of simulations run is limited. So this primary purpose of this final step is to make small corrections, not perform all-at-once optimization (which would likely require thousands of simulations). It also provides an opportunity to rebalance the contributions from multiple inputs in regions where there is a high degree of parameter inter-dependence. However, if the user is certain that they want to perform all-at-once optimization (which would likely require many more simulations), they could set the number of simulations for all steps except the last one to 0, and specify a very large number of simulations for the final step.

For each optimization step, HNN uses the COBYLA optimization algorithm (Powell, 1994), which supports bound constraints as defined by the user for each parameter. We have found COBYLA converges at a local minimum faster than the PRAXIS algorithm (Brent, 1973) as implemented in NEURON’s multiple run fitter.

## Acknowledgments

Research supported by NIH NIBIB BRAIN Award 5-R01-EB022889-02, NIH NIBIB BRAIN Award Supplement R01EB022889-02S1, NIH NIDCD 5-R01DC012947-07, ARO W911NF-19-1-0402.

## Supplementary Materials

### Sensitivity Analyses of ERP Simulations

To reduce the computational demands of performing model optimization of HNN ERP simulations, we used variance based sensitivity analysis to identify parameters that were less significant to the simulated dipole waveform. As discussed above, HNN’s model optimization feature focuses on estimating parameters of the exogenous driving inputs. Of those parameters, we sought to find ones that did not vary model output significantly and were not necessary to include in parameter estimation.

The method of variance based sensitivity analysis through Monte Carlo estimation (Sobol, 2001) provides Sobol sensitivity indices that can be used to explain the relative contribution of individual parameters on model variance. The total Sobol sensitivity index for each parameter serves as a measure that represents that parameter’s contribution to the variance, and also the contributions resulting from interactions with other parameters being varied (Homma & Saltelli, 1996). So a parameter with a low total Sobol sensitivity index can be characterized as an overall insignificant contributor to variance and can be fixed at its default value during model optimization.

We used Uncertainpy (Tennøe, Halnes, & Einevoll, 2018) to perform sensitivity analyses of parameters belonging to the exogenous driving inputs in the perceptual threshold-level (“yes_trial_SI_ERP_all_avg.txt”) and suprathreshold-level (“ERPYesSupraT.txt”) evoked response examples provided with HNN and described in the Results (Figure 4 and Figure 10). These analyses were performed using a modified simulation interface to run the simulations in parallel on a high-performance computing cluster, which is not currently included with HNN distribution. The results from our sensitivity analyses are shown in Figure 4 Supplementary Figure 1, Figure 10 Supplementary Figure 2, and Figure 10 Supplementary Table 1. Each analysis consisted of varying all parameters (except input timing standard deviation) of a driving input over 55,000 simulations using a quasi-Monte Carlo method that sampled from parameter distributions we specified. The input time distribution was defined as a normal distribution with mean and standard deviation from the default parameter file. The values for various synaptic weights were chosen from a uniform distribution ranging from the default value plus or minus 500%. For synaptic weight parameters with a default value of 0, the uniform distribution ranged from 0 to 1.0.

Because the calculation of the total Sobol sensitivity index is carried out at each point in time, and is relative to the variance at that point, total Sobol indices at different time points cannot be directly compared, and an average across the entire simulation is not appropriate. We were interested in comparing the contribution of each parameter to the dipole waveform (in units of nAM) across the whole simulation, so we computed a weighted total Sobol index at each point in time (weighted by a scaled std. deviation ranging from 0 to 1). The plots in Figure 4 Supplementary Figure 1 and Figure 10 Supplementary Figure 2 show weighted total Sobol indices for each parameter over the duration of the simulation. Supplementary Table 1 ranks the parameters with the greatest contribution to model output using the arithmetic means of weighted total Sobol indices across the entire simulation, for each driving input.

The results from our sensitivity analyses of sensory evoked response examples illustrate that there are several candidate parameters for excluding from model optimization. Not surprisingly input timing is an important parameter to optimize. In most cases NMDA weights have a greater contribution than AMPA, as do connections to Layer 5 neurons compared to Layer 2/3 neurons.

## Supplementary Figures and Tables

**Figure 10 Supplementary Figure 1:**
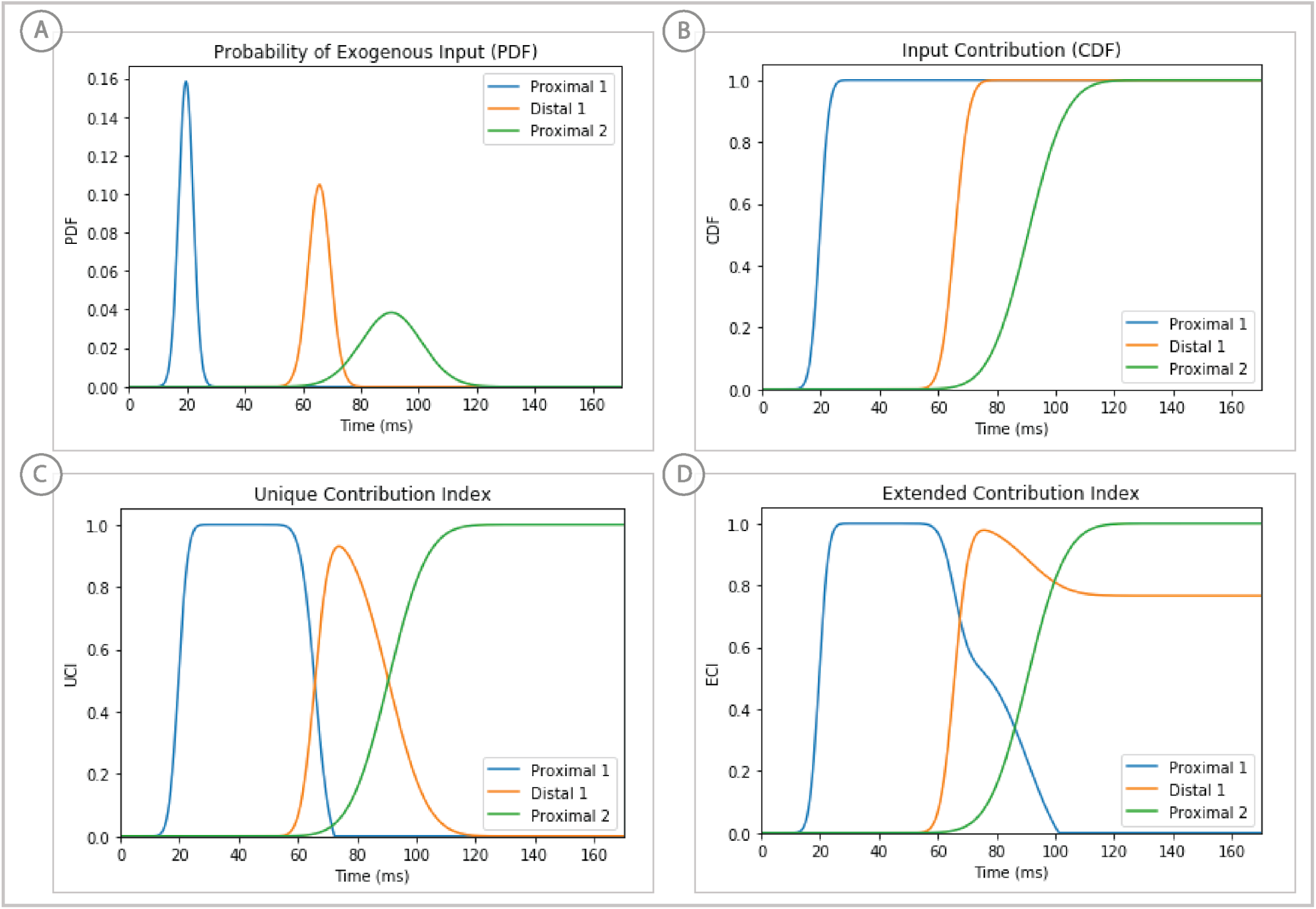
Weighted scoring of stepwise optimization procedure for suprathreshold level evoked response example, see text for details. **(A)** Probability distribution of input timing for each input. **(B)** Corresponding cumulative distribution as a contribution index. **(C)** Unique Contribution Index that is reduced by subsequent input’s CDFs. **(D)** Extended Contribution Index in which the subtracted CDFs are subject to a decay factor.

**Figure 10 Supplementary Figure 2:**
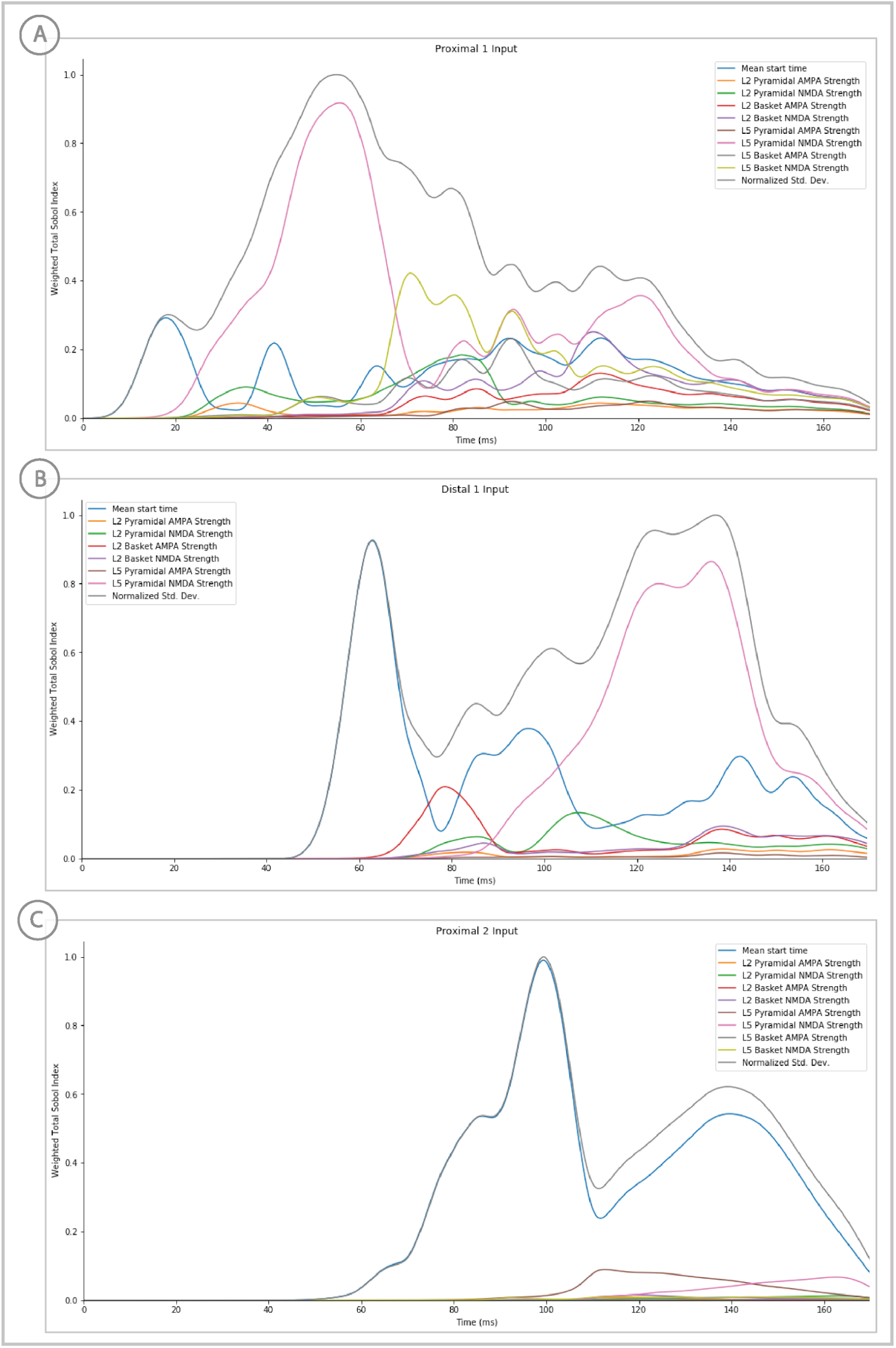
Sensitivity analysis results of the suprathreshold level evoked response example showing the relative contribution of each input’s parameters on variance. Total Sobol indices at each point have been weighted by the std. deviation scaled from 0 to 1.

**Figure 4 Supplementary Figure 1:**
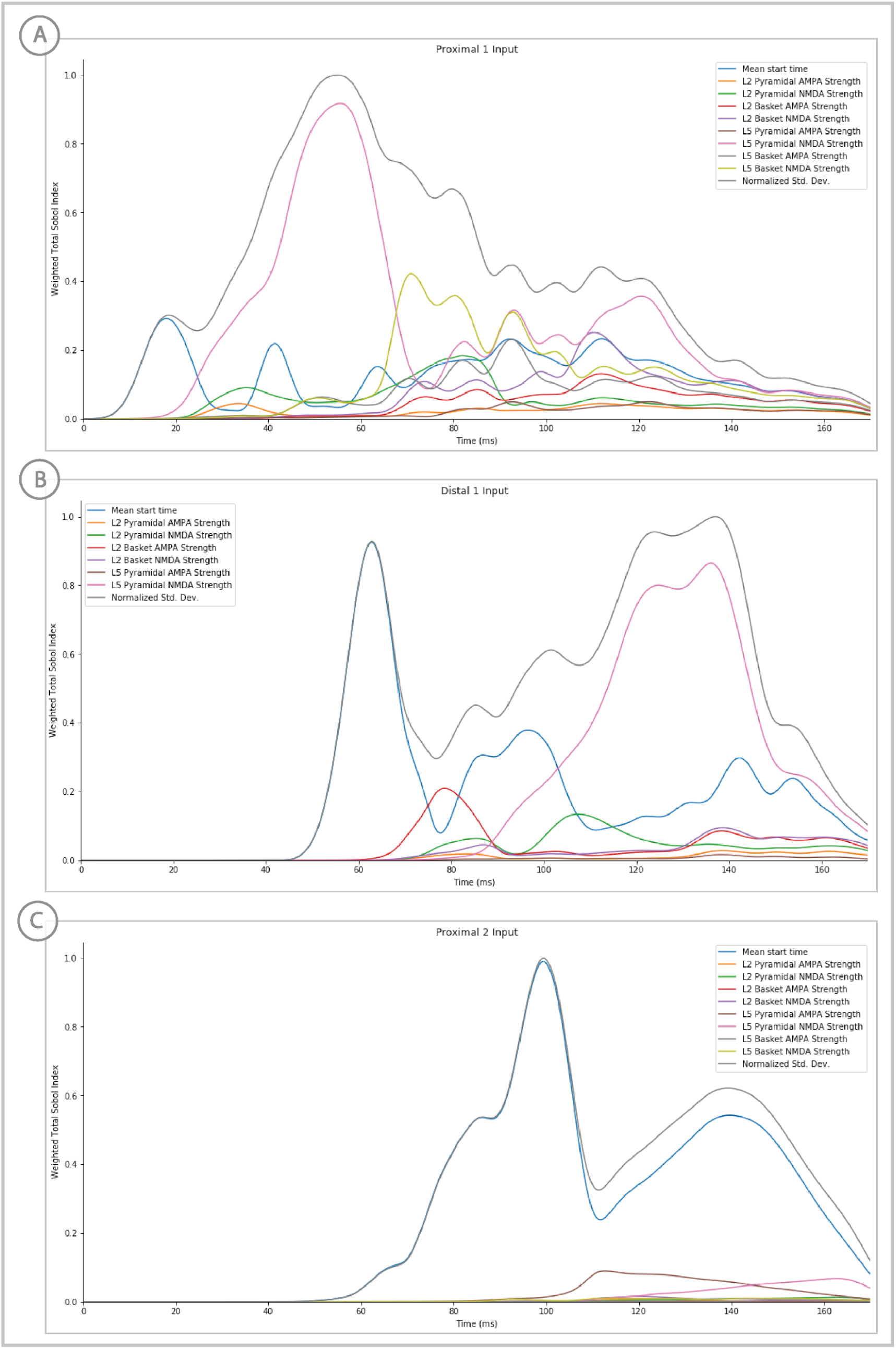
Sensitivity analysis results of the perceptual threshold level evoked response example showing the relative contribution of each input’s parameters on variance. Total Sobol indices at each point have been weighted by the std. deviation scaled from 0 to 1.

**Supplementary Table 1:** Summary of weighted total Sobol sensitivity index values averaged across the entire simulation for each exogenous driving input in two sensory evoked response models: suprathreshold (Figure 10) and 50% detected (Figure 4). The ranking of an input’s parameters remains similar between models.

| Driving Input | Parameter name | ERP Model -- 50% detected |  | ERP Model -- 100% detected |  |
| --- | --- | --- | --- | --- | --- |
|  |  | Average Weighted Total Sobol Index | Rank | Average Weighted Total Sobol Index | Rank |
| Proximal Input 1 | Mean start time | 0.2326 | 3 | 0.3059 | 2 |
|  | L2 Pyramidal NMDA Strength | 0.1458 | 5 | 0.1349 | 6 |
|  | L2 Pyramidal AMPA Strength | 0.0111 | 9 | 0.0519 | 8 |
|  | L5 Pyramidal NMDA Strength | 0.5952 | 1 | 0.6243 | 1 |
|  | L5 Pyramidal AMPA Strength | 0.0289 | 8 | 0.0449 | 9 |
|  | L2 Basket NMDA Strength | 0.0982 | 6 | 0.1702 | 4 |
|  | L2 Basket AMPA Strength | 0.0486 | 7 | 0.1055 | 7 |
|  | L5 Basket NMDA Strength | 0.2998 | 2 | 0.2748 | 3 |
|  | L5 Basket AMPA Strength | 0.1561 | 4 | 0.1672 | 5 |
| Distal Input 1 | Mean start time | 0.4442 | 2 | 0.4545 | 2 |
|  | L2 Pyramidal NMDA Strength | 0.0681 | 3 | 0.0781 | 4 |
|  | L2 Pyramidal AMPA Strength | 0.0000 | 7 | 0.0202 | 6 |
|  | L5 Pyramidal NMDA Strength | 0.5631 | 1 | 0.5163 | 1 |
|  | L5 Pyramidal AMPA Strength | 0.0292 | 5 | 0.0099 | 7 |
|  | L2 Basket NMDA Strength | 0.0432 | 4 | 0.0615 | 5 |
|  | L2 Basket AMPA Strength | 0.0149 | 6 | 0.0924 | 3 |
| Proximal Input 2 | Mean start time | 0.8424 | 1 | 0.9018 | 1 |
|  | L2 Pyramidal NMDA Strength | 0.0068 | 6 | 0.0101 | 6 |
|  | L2 Pyramidal AMPA Strength | 0.0121 | 5 | 0.0045 | 7 |
|  | L5 Pyramidal NMDA Strength | 0.1667 | 2 | 0.0475 | 3 |
|  | L5 Pyramidal AMPA Strength | 0.0514 | 3 | 0.0726 | 2 |
|  | L2 Basket NMDA Strength | 0.0041 | 7 | 0.0115 | 5 |
|  | L2 Basket AMPA Strength | 0.0000 | 9 | 0.0001 | 9 |
|  | L5 Basket NMDA Strength | 0.0223 | 4 | 0.0128 | 4 |
|  | L5 Basket AMPA Strength | 0.0032 | 8 | 0.0026 | 8 |

